# Aging modifies microstructure and material properties of mineralized cartilage and subchondral bone in the murine knee

**DOI:** 10.64898/2026.04.02.716015

**Authors:** Laura Müller, Stéphane Blouin, Edoardo Pedrinazzi, G. Harry van Lenthe, Alexandre Hego, Richard Weinkamer, Markus A. Hartmann, Davide Ruffoni

## Abstract

The osteochondral junction is a specialized region ensuring the biomechanical and biological integration of the unmineralized articular cartilage with the subchondral bone through an intermediate layer of mineralized cartilage. This location is of clinical relevance, being the target of osteoarthritis. While aging is considered a risk factor for osteoarthritis, the interplay between microstructural and material changes during aging and predisposing to joint degeneration is not fully clear. This is especially true for mineralized cartilage, which remains understudied despite its critical role in load transfer from unmineralized articular cartilage to bone. We investigate age-related alterations of mineralized cartilage and subchondral bone in rat tibiae of adult and aged animals using a multimodal, high-resolution, correlative analysis. Our approach includes micro-computed tomography to measure microstructural features, second harmonic generation imaging to visualize collagen organization, quantitative backscattered electron imaging to map local mineral content, and nanoindentation to obtain mechanical properties. Mineralized cartilage and subchondral bone exhibited distinct age-related modifications. At the architectural level, the subchondral plate thickened and the trabecular network became coarser, those changes being different from those observed in the metaphysis. At the tissue level, mineralized cartilage was less mineralized than bone but exhibits a greater relative increase of mineral content with age, underlying differences in mineralization. A central observation is that aging led to an abrupt transition in mineral content and mechanical properties across the interface between unmineralized and mineralized cartilage, with a conceivable impact on stress localization. Overall, these changes may alter load transfer and contribute to age-related joint degeneration.

## 1. Introduction

The bone-cartilage interface, or osteochondral junction, is a complex region ensuring the biomechanical and biological integration of two tissues, bone and cartilage, which have dissimilar properties and functions [1]. This region is not only fundamental for joint functioning but also of high clinical relevance, being a target of osteoarthritis (OA), the most prevalent joint disease, with primary risk factors being aging and injury [2–4]. The bone-cartilage interface is also interesting from a materials engineering standpoint, as it withstands high dynamic stresses for several decades. A challenging task, which is further intensified by the requirement to connect, within a zone that is only a few hundred micrometers in width, a soft tissue - hyaline cartilage, having an elastic modulus on the order of tens of megapascal - with a much stiffer one - subchondral bone, which can reach stiffness values up to 15-20 GPa. Biomaterial interfaces are prone to stress concentration and damage [5,6]; yet, nature has found several solutions to mitigate, dissipate, and withstand such high stresses [7–10].

Analogous to the tendon-to-bone attachment at the enthesis [11–14], the connection between unmineralized articular cartilage and bone is mediated by mineralized cartilage. Despite being a key element of the osteochondral junction, located on the force trajectory from cartilage to bone, mineralized cartilage remains relatively understudied. Contributing factors include its very small size, its difficult-to-access location, and the historical “prejudice” considering mineralized cartilage as a mere by-product of endochondral ossification, rather than a functional, mechanoresponsive, and metabolically active tissue [15]. This perspective is now being reconsidered due to growing evidence of biomechanical [8,16] and mechanobiological [17] interactions between mineralized cartilage and surrounding tissues. At the material level, both mineralized cartilage and bone use nanometer-sized mineral particles to reinforce a fibrous collagen network [18,19], which is based on collagen Type II in cartilage and collagen Type I in bone. The two tissues, however, possess different mineral-mechanics relationships: in humans, for the same amount of mineral, subchondral bone shows higher stiffness and hardness than mineralized cartilage [20]. The opposite behavior is observed in mineralized (fibro)cartilage in rats [12]. These differences are not fully understood, highlighting that a comprehensive picture of the composition–structure-mechanics relationship in mineralized cartilage is still missing. Unfortunately, it is even less clear how aging and disease influence this relationship. Structurally, mineralized cartilage is bordered by two distinct interfaces [15,21]: the tidemark denotes the transition between non-mineralized and mineralized cartilage, and the osteochondral cement line is found between mineralized cartilage and subchondral bone. Type II collagen fibrils in hyaline cartilage are thought to cross the tidemark at approximately right angles, helping to resist shear stresses [18]. The fibrils extend into mineralized cartilage, where they get reinforced by mineral particles and “glue” to neighboring Type II collagen fibrils. A steep increase in mineral content and mechanical properties (e.g. elastic modulus and hardness) is usually observed across the tidemark over a narrow region of only a few micrometers in width [9,20,22,23]. At the same location, nanoscale modifications in mineral morphology, density, and crystallinity have been reported and assumed to improve energy dissipation [7,10]. The anchoring of mineralized cartilage to subchondral bone is ensured by a highly intricate and interlocked interface [21]. The possible contribution of direct bonding between collagen fibrils across the osteochondral cement line is debated, as fibrils seem to have different orientations at the opposite side of the interface [18,24]. Yet, there is some evidence that Type I and Type II collagen fibrils coexist at the cement line, providing an option for bonding between the two tissues [25].

With respect to its biological functions, mineralized cartilage is nowadays considered an active tissue, showing loading, injury, aging, and disease-related changes [15,21,26,27]. Once endochondral ossification is completed, two dynamic processes are responsible for modifying mineralized cartilage. Firstly, a slow progression of the mineralization front—from mineralized towards unmineralized cartilage—leads to tidemark advancement and multiplication. An important unresolved question is how aging modifies the mineralization front: as it advances into articular cartilage in aged joints, is the corresponding mineralization gradient altered? If so, this could have consequences for energy dissipation. The second process, occurring on the opposite side of mineralized cartilage, is its resorption and subsequent replacement with subchondral bone. These two biological processes influence the thickness and morphology of mineralized cartilage, but their impact on material-level properties remains insufficiently characterized. Similar to bone [28,29], mineralized cartilage shows a heterogeneous degree of mineralization [9,27,30,31]. How the degree of mineralization of this tissue evolves with age is unclear, although it is a fundamental aspect to explain changes in the local biomechanical properties [9,12,20].

Motivated by those unanswered questions, we follow a multimodal, correlative, high-resolution approach to investigate age-related changes at the osteochondral junction, focusing on mineralized cartilage and the underlying subchondral bone. Specifically, we combine, on the same samples and locations, micro-computed tomography (micro-CT) to assess microstructure, quantitative backscattered electron imaging (qBEI) to evaluate mineral content and porosity, second harmonic generation (SHG) imaging to visualize collagen organization, and nanoindentation (nIND) to probe local biomechanical properties. To study the effect of aging in a controlled setting and independently from OA, we characterize the osteochondral region in rat knees. The similarity between rat and human knee biomechanics is one reason why rats are frequently used as pre-clinical models in orthopedic and cartilage research. Our central aim is to clarify how the mineralized tissues of the osteochondral junction change with age, a key step toward identifying mechanisms driving joint degeneration.

## 2. Materials and Methods

### 2.1. Samples

We considered tibiae from 3- and 15-month-old male Wistar rats (n=10 in both groups). Animals were obtained through an organ donation program at the Liège University Hospital (CHU), conducted under the approval of the Animal Ethics Committee of the University of Liège (IACUC-22-2416). All animals were housed in ventilated cages (6–7 rats per cage) placed in an environmentally controlled room at a 12-hour light/dark cycle, with free access to a standard diet and water. At the time of sacrifice (by pentobarbital), adult and aged rats had a mean body weight of 459 g and 681 g, respectively (Supplementary Information, Table S1). Tibiae were carefully extracted immediately after sacrifice using a scalpel, taking care to preserve the thin layer of articular cartilage. Following harvesting, all samples were immersed in 70% ethanol for preservation until further processing.

### 2.2. Microcomputed tomography (micro-CT) and image processing

To obtain microstructural information, the proximal tibiae were scanned using micro-computed tomography (micro-CT, Skyscan 1272, Bruker) at a nominal isotropic voxel size of 10 µm. Relevant scanning parameters include a tube voltage of 90 kVp, a current of 110 µA, and a rotation step of 0.4° over 180°. The exposure time was 2300 ms, and a frame averaging of 4 was used to reduce noise. To minimize beam hardening artifacts, a 0.5 mm-thick aluminum and a 0.038 mm-thick copper filter were used. Image reconstruction was performed using a filtered back-projection algorithm, including ring artifact reduction and beam hardening correction, implemented in the software of the scanner (NRecon, Bruker). After scanning, an image processing workflow was implemented to investigate bone morphology in the epiphysis (including the subchondral region) and in the metaphysis (used as a reference). To reduce high-frequency noise, a three-dimensional Gaussian filter (square kernel size of 1.2) was applied prior to binarization using a global threshold approach based on the Otsu method [32]. Both operations were performed with the software CTAn (Bruker). The fibula was digitally removed using the same software. After aligning all bones along the same orientation, the epiphysis and the metaphysis were identified by manual contouring. In both regions, trabecular bone was then isolated from the cortical shell using an automatic approach based on standard morphological operators as implemented in CTAn (details on bone alignment and trabecular segmentation can be found in the Supplementary Information, Note 1 and Fig. S1). In the epiphysis, trabecular bone was further subdivided into medial and lateral compartments using the intercondylar eminence as an anatomical landmark. Trabecular bone morphology, including bone volume fraction (BV/TV), trabecular thickness (Tb.Th) and trabecular separation (Tb.Sp) was quantified with CTAn following standard guidelines [33]. The subchondral plate (SCP) was also isolated and analyzed in terms of thickness and curvature. Due to the resolution of the micro-CT, the SP encompasses both subchondral cortical bone and mineralized cartilage. The subchondral plate thickness (SCP.Th) was obtained in two steps: firstly, the plate was slightly smoothed to reduce the roughness, and then the nearest neighbor algorithm [34], computing the shortest distance between the upper and the bottom surface, was used (details on plate extraction and analysis are given in the Supplementary Information, Note 2). The same approach was followed to measure the whole height of the epiphysis, defined as the mean distance from the plateau (i.e. the top articular surface that articulates with the femoral condyles) to the growth plate and restricting the analysis to the region beneath the plateau. The surface curvature of the subchondral plate was computed in MATLAB (MathWorks) by fitting the extracted top surface with a fifth-degree polynomial, using a two-variable polynomial surface (poly55 in MATLAB) and linear least squares. The fitted surface, 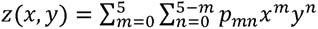, was then used to calculate the local Gaussian curvatur (SCP.K) [35]. In the metaphysis, the average thickness of the cortical shell, Ct.Th, was computed using the sphere fitting method [36].

### 2.3. Sample preparation for material analysis

After micro-CT, samples were prepared for material characterization, including quantitative backscattered electron imaging (qBEI), nanoindentation (nIND), and second harmonic generation imaging (SHG). Specifically, six samples per group were analyzed using qBEI; a subset of three samples per group was then further characterized using nIND and SHG. The three techniques were performed on the same surfaces and locations, enabling a correlative analysis. The samples were dehydrated and defatted with a series of ascending grades of ethanol, acetone, and methylmethacrylate (MMA) solutions, before embedding in polymethylmethacrylate (PMMA), following a well-established protocol [37]. The embedded tibiae were then cut along the frontal plane using a diamond wire saw (DWS.100, Diamond WireTec). The exposed surfaces were ground with sandpaper until the two knee condyles (and corresponding plateaus) as well as the intercondylar valley were clearly visible under an optical microscope, and had approximately the same dimensions, ensuring consistency among all samples. These features are highlighted in Fig. 1c as an example. A final polishing step using a diamond suspension with a grain size of 1 μm (Logitech PM5) was performed. Grinding and polishing were done under glycol irrigation to avoid microcrack formation due to water [38].

**Figure 1:**
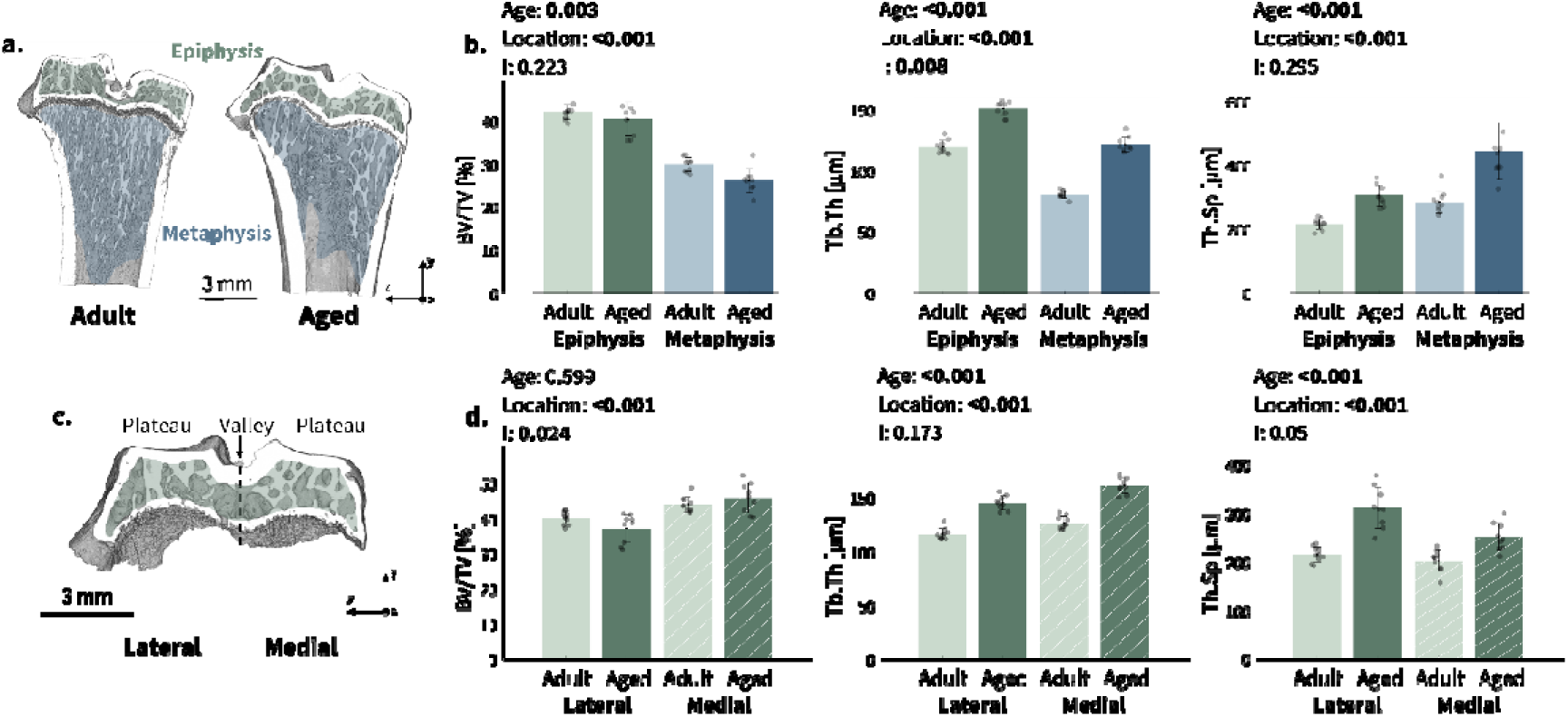
Microstructural analysis of trabecular bone. Micro-CT images sectioned along a frontal plane showing trabecular microstructure in (a) epiphysis and metaphysis of adult and aged rats, and in (c) epiphyseal regions of an aged animal. Trabecular bone parameters (BV/TV: bone volume fraction; Tb.Th: trabecular thickness; Tb.Sp: trabecular spacing) computed in (b) metaphysis and epiphysis and in (d) lateral and medial compartments of the epiphysis. Data shown as mean (histogram height) with bars representing one standard deviation. Individual samples are reported as scattered points. Results of two-way ANOVA are reported considering age, location and their interaction (I).

### 2.4. Quantitative backscattered electrons imaging (qBEI)

Mineral content was quantified with qBEI, a well-established 2-dimensional technique where the surface of the sample is scanned with an electron beam and the intensity of the electrons that are back-scattered from a thin surface layer (about 1-2 µm in thickness) is detected [28,37,39]. In bone, the backscattered signal is essentially dominated by the local calcium concentration [37]. Prior to qBEI measurements, a thin carbon layer was deposited (AGAR SEM carbon coater) to ensure a conductive surface and to prevent charging effects. The measurements were done using a Field Emission Scanning Electron Microscope (FESEM, Supra40, Zeiss) operating at 20 kV, 270 to 320 pA sample current, with a working distance of 10 mm and at a magnification of x130, yielding a nominal isotropic pixel size of 0.88 μm [39]. Several images of 1024 × 768 pixels were acquired with a spatial overlap of 5% and stitched together, such that the entire frontal section of the proximal tibia could be imaged. A daily calibration of the FESEM was performed according to a previously established procedure using carbon and aluminum standards [37]. This allowed the conversion of gray levels (GL) to calcium weight percentage (wt % Ca) using the following linear equation, with a GL of 25 corresponding to osteoid (0 wt % Ca) and of 255 to pure hydroxyapatite (39.86 wt % Ca):

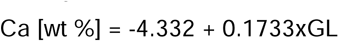

Evaluation of qBEI images started with a manual segmentation (done in CTAn) of mineralized cartilage and subchondral bone following the osteochondral cement line (shown in Fig. 3 as dashed line), which delineates the interface between the two tissues. A minimum threshold of 5.2 wt % Ca was used to exclude signals from embedding resin and non-mineralized tissues. To minimize partial volume effects [28], a 2-pixel erosion was performed. Examination of the qBEI maps highlighted the presence of some chondrocyte lacunae partially or completely filled with minerals, as well as of a few protrusions of high mineral content extending from mineralized into non-mineralized cartilage (Fig. 3). Both features, which will be addressed later, were manually discarded from the quantification of the qBEI dataset. To evaluate the mineral content in approximately the same locations across the different samples, two regions of interest were defined (white boxes in Fig. 4), spanning the whole medial and lateral tibial plateau and extending into subchondral bone up to a depth of 40% of the total epiphyseal height (corresponding to approximately 600 µm), measured at the center of each plateau. Based on the mineral content of these regions, frequency distributions of the mineral content (normalized to unit area) were computed for bone and mineralized cartilage. The distributions, informing on the heterogeneous mineral content of the tissues [29,40], were characterized by three parameters [28]: Ca_Mean_ (average Ca concentration), Ca_Peak_ (most frequent Ca concentration), and Ca_Width_ (full width at half maximum). The mineralized cartilage layer was also used to estimate the thickness of this tissue (which was not distinguishable from the subchondral bone in micro-CT), using the sphere fitting technique [36]. Finally, the spatial variation in mineral content across the tidemark as well as inside mineralized cartilage was characterized with two approaches. Locally, on the original qBEI images, we computed profile plots along 15 probe lines (3-pixel width and 40-pixel long) perpendicular to the tidemark on each plateau and sample. A sigmoid function was fitted to the resulting profiles, and the transition width, defined as the difference between 0.5% and 99.5% height of the sigmoid, provided a quantitative estimation of the steepness of the mineralization front [12]. A second analysis provided a more global characterization of the mineral content in mineralized cartilage with respect to the distance to the tidemark. Specifically, we assessed the probability of finding, at a given distance from the tidemark, a pixel with a specific mineral content. The analysis is based on the distance transform function computed to the tidemark, assigning positive distance values when moving into mineralized cartilage. At each distance, discretized with a bin size of 0.88 µm, the frequency distribution of mineral content was calculated. The distribution was normalized relative to the sum of all pixels at each distance, allowing to correct for possible bias due to a different number of pixels at different distances to the tidemark [41].

**Figure 2:**
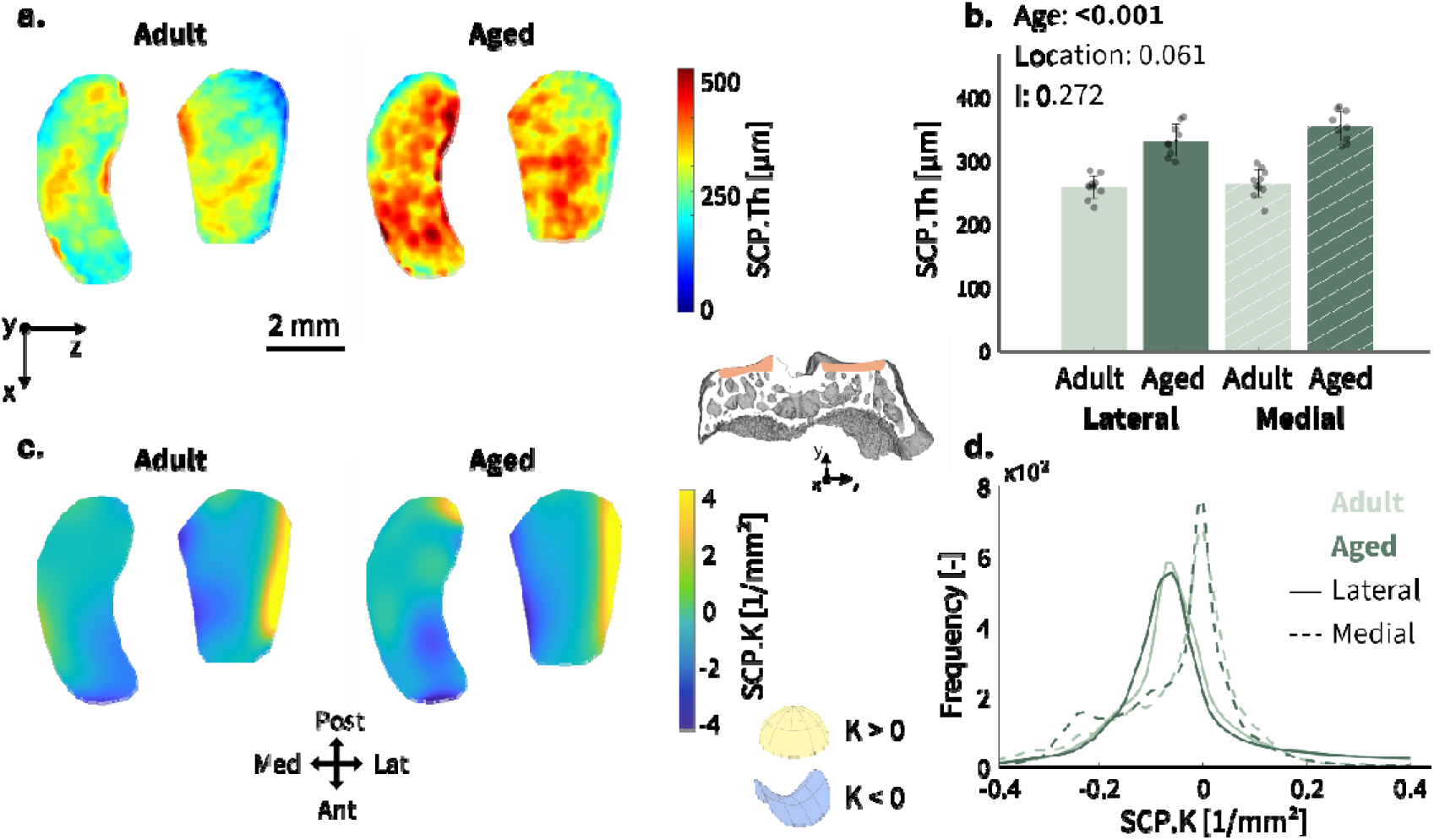
Microstructural analysis of the subchondral plate. Representative maps of (a) subchondral plate thickness (SCP.Th) and (c) local Gaussian curvature (SCP.K) projected on the top surface of the tibia for an adult and aged rat. (b) Subchondral plate thickness in adult and aged settings, considering lateral and medial plateaus. Data shown as mean (histogram height) with error bars representing one standard deviation. Individual samples are shown as scattered points. Results of two-way ANOVA are reported considering age, location and their interaction (I). (d) Frequency distributions (normalized to unit area) of Gaussian curvature in adult and aged rats for lateral and medial plateaus. Also shown in the figure is a micro-CT image of the subchondral region (frontal view), highlighting the two plateaus.

**Figure 3:**
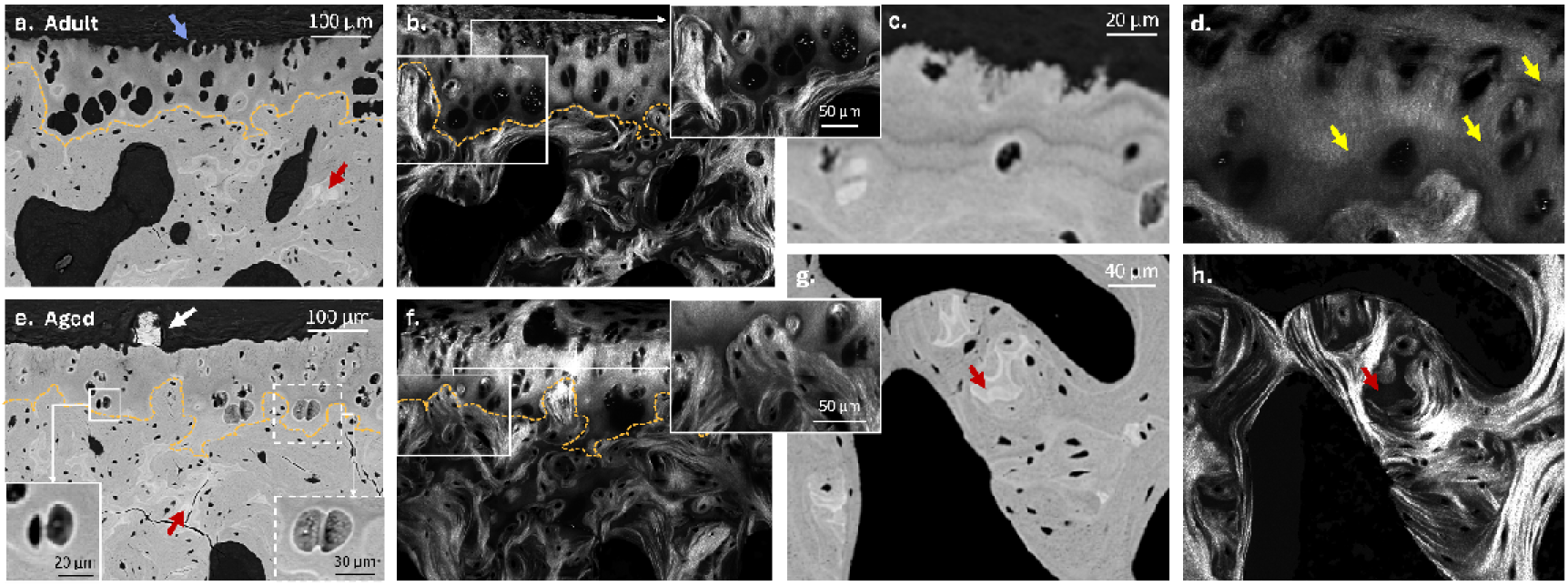
Microporosity and fiber organization. qBEI maps of a representative (a) adult and (e) aged sample (frontal section) showing mineralized cartilage and subchondral bone. The two insets in (e) offer a magnified view on chondrocyte lacunae bordered by a thin hypermineralized layer (full box) and mineral-filled chondrocyte lacunae (dotted box). In (a, e), the following features are highlighted: an arrested chondrocyte (blue arrow), a highly mineralized protrusion invading articular cartilage (white arrow), and highly mineralized cartilaginous islands (red arrows). In (b, f) the same regions are imaged with SHG. The orange dashed line denotes the osteochondral cement line. The two insets show magnified views at the osteochondral cement line where no SHG signal is detected. (c) qBEI map highlighting former tidemarks in mineralized cartilage of an aged sample and (d) corresponding SHG image showing variations in intensity colocalizing with the tidemarks (yellow arrows). (g) qBEI and (h) corresponding SHG image of subchondral trabecular bone, underlying the presence of cartilaginous inclusions and lamellar patterns.

**Figure 4:**
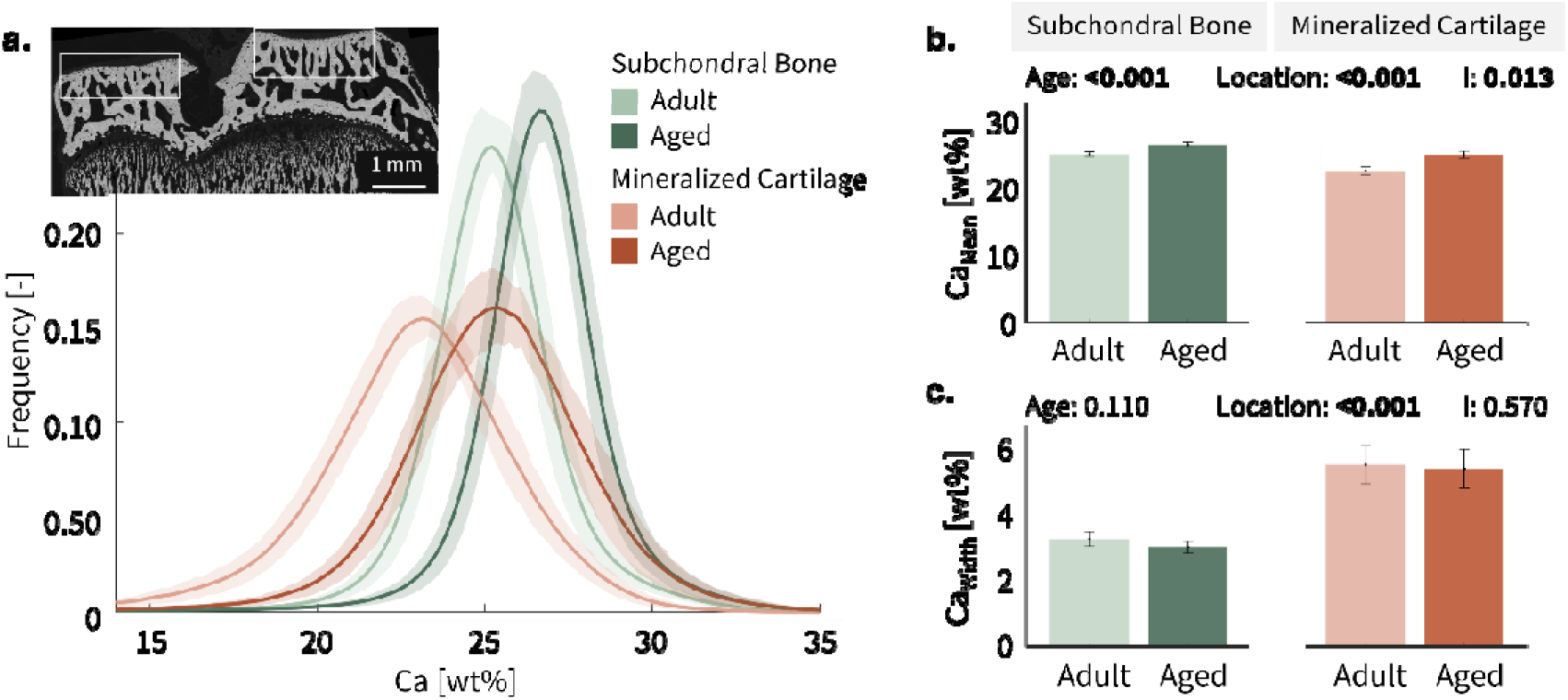
Evaluation of the spatial heterogeneity of the mineral content. (a) Frequency distributions of calcium content in subchondral bone (green) and mineralized cartilage (red) shown as smoothed splines. Lighter colors indicate adult samples and darker colors aged ones. Full lines indicate mean values and shaded areas one standard deviation interval. The inset shows a representative qBEI (frontal section), and the white boxes enclose the regions used to compute the frequency distributions (separating mineralized cartilage from bone, see also Fig. 3). Comparison of (b) mean mineral content (Ca_Mean_) and (c) mineralization heterogeneity (Ca_Width_) in subchondral bone and mineralized cartilage between adult and aged rats. Data shown as mean (histogram height) with error bars representing one standard deviation. Results of two-way ANOVA are reported considering age, location and their interaction (I).

### 2.5. Depth-sensing nanoindentation (nIND)

The local mechanical behavior of the tissues was probed with depth-sensing nanoindentation (nIND). Before testing, the carbon coating was removed using a diamond polishing paste. nIND was done using a TI950 TriboIndenter (Bruker) equipped with a Berkovich diamond probe (with the probe area calibrated on fused quartz). To ensure a comparable penetration depth across the tidemark from non-mineralized to mineralized cartilage, a displacement-controlled trapezoidal load function was used (10s loading, 15s holding, and 10s unloading), with a maximum penetration depth of 150 nm. The average surface roughness around the indent, measured by scanning 25 μm x 25 μm regions with the tip of the nanoindenter at 2 μN contact force, was 4.89 ± 3.18 nm. The lateral spacing between adjacent indents was set to 1.5 µm, allowing for a high-resolution 2D mechanical mapping, while avoiding overlapping of the inelastic deformation field [9]. Indentation modulus and hardness were extracted from the load–displacement curves using the Oliver–Pharr method [42]. Several nanoindentation grids were performed across the tidemark on both plateaus, with some grids extending until subchondral bone. Grid locations were selected based on the qBEI maps, avoiding regions with too many cell lacunae, pores, and cracks. A total of approximately 5000 and 4600 indents were performed on samples from 3- and 15-month-old animals, respectively. The nanoindentation grids were then manually overlaid onto the qBEI images using recognizable features (tidemark, pores), enabling the distinction between indents located in mineralized cartilage and subchondral bone, as well as the calculation of local mineral content in the same regions probed by nanoindentation. In the quantification of average mechanical properties within mineralized cartilage and bone, all points with indentation modulus below 10DGPa, likely compromised by the presence of pores/cracks beneath the surface and therefore not visible with qBEI, were excluded.

### 2.6. Second harmonic generation (SHG) imaging

As a final characterization, SHG was performed on the same samples used for nIND. The osteochondral junction was imaged with a confocal microscope (NIKON A1R MP+ multiphoton) equipped with a 40x oil immersion objective (numerical aperture of 1.15). The microscope was operated with a voltage of 110V, a laser power of 0.16 W, and a wavelength of 840 nm, producing images with a nominal isotropic pixel size of 0.3 µm and a field of view of 1024 ×1024 pixels with a spatial overlap of 10% stitched together such that central region of both tibial plateaus was imaged. This technique allows characterization of the spatial organization of collagen fibrils across the osteochondral junction, with the SHG signal being particularly strong when collagen fibrils are densely packed and aligned parallel to the imaging plane, whereas no signal is detected when they are oriented perpendicularly to this plane [43–45].

### 2.7. Statistical methods

To compare the effect of age and location on microstructural and mineralization parameters we used a two-way repeated-measures analysis of variance (ANOVA). Data normality was verified using the Shapiro–Wilk test. In case of significant interaction between factors, Holm–Sidak post hoc comparisons were performed. When assessing only the effect of age, a Student’s t-test was performed, with normality checked by a Shapiro–Wilk test. Significance level was set to 0.05. All statistical analyses were conducted using SigmaPlot 12.5 (Systat Software).

## 3. Results

From a qualitative observation of the micro-CT images, we noticed that one sample of the aged group showed clear alterations in the shape of the medial plateau as well as in the microstructure of the subchondral trabecular bone. The qBEI maps also revealed a bony layer above mineralized cartilage in the lateral plateau, terminating in an osteophyte (both micro-CT and qBEI images are in the Supplementary Information, Fig. S2). Since these modifications are potential indicators of joint degeneration and were not observed in the other bones, the sample was considered an outlier and excluded.

### 3.1. Microstructural changes: trabecular bone

A two-way ANOVA with age and anatomical location (epi- and metaphysis) as factors showed significant effects on all the considered trabecular parameters (BV/TV, Tb.Th and Tb.Sp), while the interaction between the two factors was only significant for Tb.Th (Fig. 1a, b). Compared to the metaphysis, the epiphysis was characterized by higher BV/TV and Tb.Sp. A post-hoc correction indicated that Tb.Th was also significantly higher in the epiphysis (in both adult and aged rats) and increased with aging by 26 % (p < 0.001) and by 51 % (p < 0.001) in epiphysis and metaphysis, respectively. Thus, aging induced bone loss and coarsening of the trabecular network, resulting in thicker and more widely spaced trabeculae.

A more detailed analysis of the epiphyseal region highlighted microstructural differences between medial and lateral compartments (Fig. 1c, d), with the latter characterized by higher BV/TV, and Tb.Th and lower Tb.Sp. With aging, Tb.Th and Tb.Sp increased in both compartments, while post-hoc test for BV/TV (which showed significant interaction between factors) suggested that this parameter decreased only in the lateral (−7 %, p = 0.049) but not in medial compartment where it stayed constant with age (p = 0.206).

### 3.2. Microstructural changes: subchondral plate

The comparison between medial and lateral compartments was further extended to the subchondral plate, characterized in terms of thickness (Fig. 2a, b) and curvature (Fig. 2c, d). While subchondral plate thickness did not depend on the specific compartment, aging led to plate sclerosis in both lateral and medial regions. Cortical bone in the metaphysis had also higher thickness in aged animals in comparison to adult ones, with the interaction between age and location (subchondral plate versus cortical bone) being statistical significance (p = 0.025, Supplementary Information, Fig. S3). A post-hoc test indicated that thickness increased more at the subchondral plate (+32 %, p < 0.001) than at the metaphyseal cortex (+24 %, p < 0.001). To evaluate whether the subchondral plates thickened mainly towards the articular cartilage (outwards) or the marrow space (inwards), we compared plate thickening to the whole height-variation of the epiphysis. As the latter did not show statistically significant variations with age (p = 0.053), while SCP.Th increased by 32% (p< 0.001), changes in subchondral plate thickness predominantly occurred inwards. Additional information is obtained by looking at the spatial distribution of local plate thickness (Fig. 2a), highlighting a heterogeneous behavior, with thicker regions localizing in the center of the plates. Such a pattern stays quite consistent across all adult samples. Moreover, regions that show higher thickness in adults also maintain relatively greater thickness with age (Supplementary Information, Fig. S4). The local Gaussian curvature of the subchondral plate (SCP.K, Fig. c) is summarized into frequency distributions (normalized to unit area), distinguishing between location and age group. In the medial region, the distributions showed a narrow peak centered around zero curvature. In contrast, the lateral plate displayed broader distributions (both in adult and aged conditions) with a peak at negative curvature values (SCP.K ͌ - 0.75).

### 3.3. Qualitative observation on tissue-level morphology, mineral content and fiber organization

Quantitative backscattered electron imaging (qBEI) was used to investigate the calcium content of mineralized cartilage and subchondral bone, as well as morphological features of the tissues. A qualitative analysis of mineralized cartilage qBEI maps (Fig. 3a, e) revealed that chondrocyte lacunae often aggregated into clusters, and larger lacunae/clusters tend to be in deep mineralized cartilage. It is also observed that adult samples exhibit higher micrometer-scale porosity than aged samples. Quantifying the relative porosity of mineralized cartilage as pore volume per tissue volume, we found values of 0.13 in adults and 0.09 in aged samples (+42 %, p < 0.01). Chondrocytes intersecting the tidemark, referred to as arrested chondrocytes [9,12,20], appeared more common in adult samples and contributed to the higher roughness of the interface between unmineralized and mineralized cartilage in comparison to aged rats (Fig. 3a, e). Another distinct feature of aged animals is that several chondrocyte lacunae were partially (or even completely) filled with mineral (Fig. 3e, green arrow) while such filled lacunae were rarely present in adult bones. Moreover, many lacunae were also bordered by a thin hypermineralized layer, again more evident in aged rats (Fig. 3e, inset). Highly mineralized protrusions (∼50–60 µm in size) extending into articular cartilage were observed at the tidemark in two aged samples (one example is given in Fig. 3e, white arrow).

The interface between mineralized cartilage and subchondral bone, delineated by the osteochondral cement line (highlighted in Fig. 3a, b, e, f), exhibited a complex morphology with interdigitations. The tortuosity of the cement line, defined as the ratio of its total length to the Euclidean distance between its start and end points, was approximately 2.08 in adult samples and 2.23 in aged samples (p = 0.403). Due to such an irregular interface, the local thickness of mineralized cartilage measured in 2D sections varied substantially, and there was not a statically significant difference between adult and ages samples (p = 0.49, Supplementary Information, Fig. S3b). Considering subchondral bone, qBEI images highlighted the presence of highly mineralized islands entrapped in bone (Fig. 3a, e, and g, red arrows). These are unremodeled cartilaginous inclusions, remnants of endochondral ossification, a known feature of rodent bone [12,46]. One last aspect worth mentioning is the presence of several cracks, which were more common in aged rats (Fig 3e and Supplementary Information, Fig. S5). As sample preparation was the same for both groups, the larger amount of cracks in aged rats, suggests differences in material properties (see next section). The SHG images provide qualitative information on the organization of collagen fibrils at the osteochondral junction (Fig. 3). In mineralized cartilage, SHG appears rather homogeneous with no evidence of lamellar organization but a difference in intensity seems to colocalize with a former tidemark (Fig. 3d). No signal is detected at the osteochondral cement line, nor within the filled chondrocyte lacunae or the highly mineralized protrusions (Fig. 3b, f, insets). Conversely, the SHG analysis of subchondral bone highlights a high heterogeneity of fibril arrangement. Several locations show lamellar patterns, for example, near marrow space and channels, intertwined with regions having a more blurred texture (Fig. 3h). Close to the osteochondral cement line, bone lamellae seem to run predominantly along this interface (Fig. 3b, inset). Areas of very weak or no SHG signals are also visible, some of them corresponding to unremodeled cartilaginous inclusions (Fig. 3h, red arrow). No major qualitative differences are observed between the two age groups, suggesting that collagen fibril organization remains largely unchanged with aging.

### 3.4. Material changes: degree of mineralization and spatial distribution of mineral content

qBEI maps were converted into frequency distributions of the mineral content (normalized to unit area), computed in mineralized cartilage and subchondral bone separately (Fig. 4a). The distribution of mineralized cartilage was shifted towards lower mineral content with respect to subchondral bone, reflected in a ∼10 % (p < 0.001) and ∼5% (p < 0.001) lower Ca_Mean_ in adult and aged animals, respectively (Fig. 4b). At the same time, mineralized cartilage always exhibited greater mineralization heterogeneity than bone, evident by a larger Ca_Width_ (Fig. 4c). With aging, both distributions moved towards higher mineral content, with the corresponding increase in Ca_Mean_ being about +6 % (p<0.001) in bone and +10 % (p<0.001) in mineralized cartilage. While shifting to higher mineral content, the frequency distribution of mineralized cartilage had practically the same shape, while the frequency distribution of subchondral bone had a higher and somewhat narrower peak. Yet, age-related changes in Ca_Width_ did not reach statistical significance and should be interpreted only as a qualitative trend (see also Supplementary Information, Table S2).

The sub-micrometer resolution of the qBEI measurements allowed a detailed characterization of the spatial increase in mineral content in mineralized cartilage (Fig. 5). Locally, the mineral content was measured along several “probe” lines perpendicular to the tidemark. In adult samples, mineral content increased progressively across the interface between unmineralized and mineralized cartilage, with a transition region of approximately 9 µm in width. In contrast, aged rats exhibited a much steeper mineralization gradient, with the width of the transition region reducing to about 4 µm (Fig. 5a, b, and Supplementary Information, Table S3). The steep gradient is further emphasized by the presence of a thin layer (around 3 µm in width) of high mineral content (28 wt %) right at the tidemark, appearing as a peak in the line profile. A more general trend is reported in Figure 5c, d and obtained by computing the frequency distributions of the mineral content as a function of the distance from the tidemark and normalized for each distance, such that the color code in the two figures indicates the probability of finding a pixel with a specific Ca value at a given distance from the tidemark. In adult animals, pixels with lower calcium content were most likely found close to the interface, while those with higher mineral content were observed furthest away (Fig. 5c). The plot also highlighted a gradual increase in mineral content near the tidemark and a much slower increase moving towards subchondral bone. In aged samples, pixels with higher mineral content were likely to be found both close to the tidemark as well as far away, as highlighted by the two dark clouds in Fig. 5d and in the inset.

**Figure 5:**
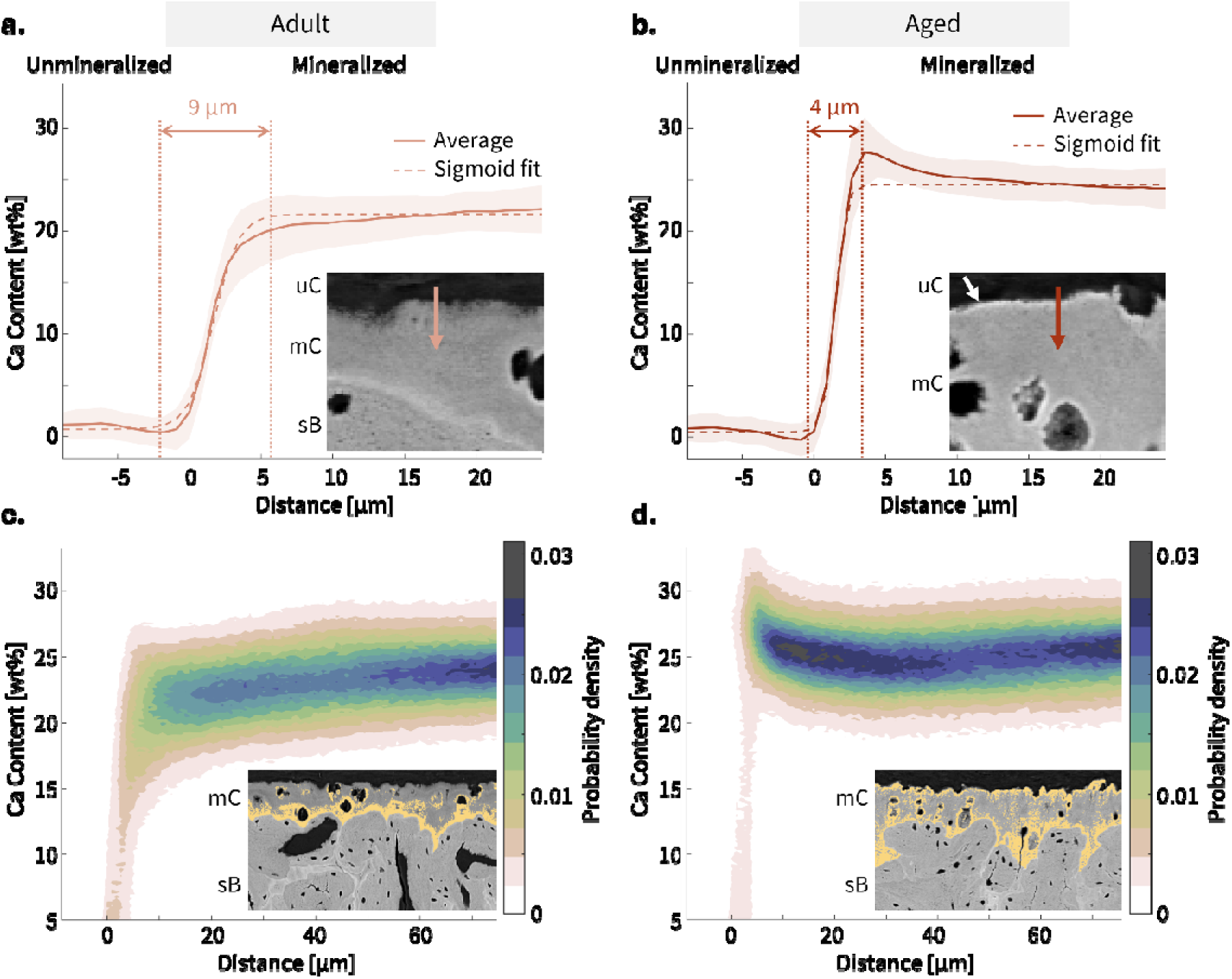
Spatial evolution of calcium content. Profile plots across the tidemark for (a) adult and (b) aged rats. The full line represents data averaged over 30 profile lines per sample, and the shaded area represents the standard deviation interval. The dashed line is a sigmoid fit (see Supplementary Information), with the corresponding transition width marked by the vertical lines. The two insets zoom on representative transition regions (uc: unmineralized cartilage, mc: mineralized cartilage, sb: subchondral bone). Mineral content probability distributions as a function of the distance from the tidemark for (c) adult and (d) aged animals. The color code indicates the probability of finding a pixel with a specific Ca value at a given distance from the tidemark. Insets show qBEI images where regions in mineralized cartilage of relatively high mineral content (>23 wt % Ca and > 26.5 wt% Ca in adult and aged samples, respectively) are highlighted in dark yellow.

### 3.3. Mechanical changes: indentation modulus and hardness

Based on the analysis of the mineral content, highlighting age-related modifications close to the tidemark, we mapped the corresponding local mechanical properties using nanoindentation on several regions, with emphasis on the unmineralized-mineralized cartilage interface (Fig. 6 and Supplementary Information, Fig. S7). Spatial variations of indentation modulus closely mirrored the behavior of calcium content at the mineralization front: adult rats had a more gradual transition in stiffness (Fig. 6a) than aged ones (Fig. 6b).

**Figure 6:**
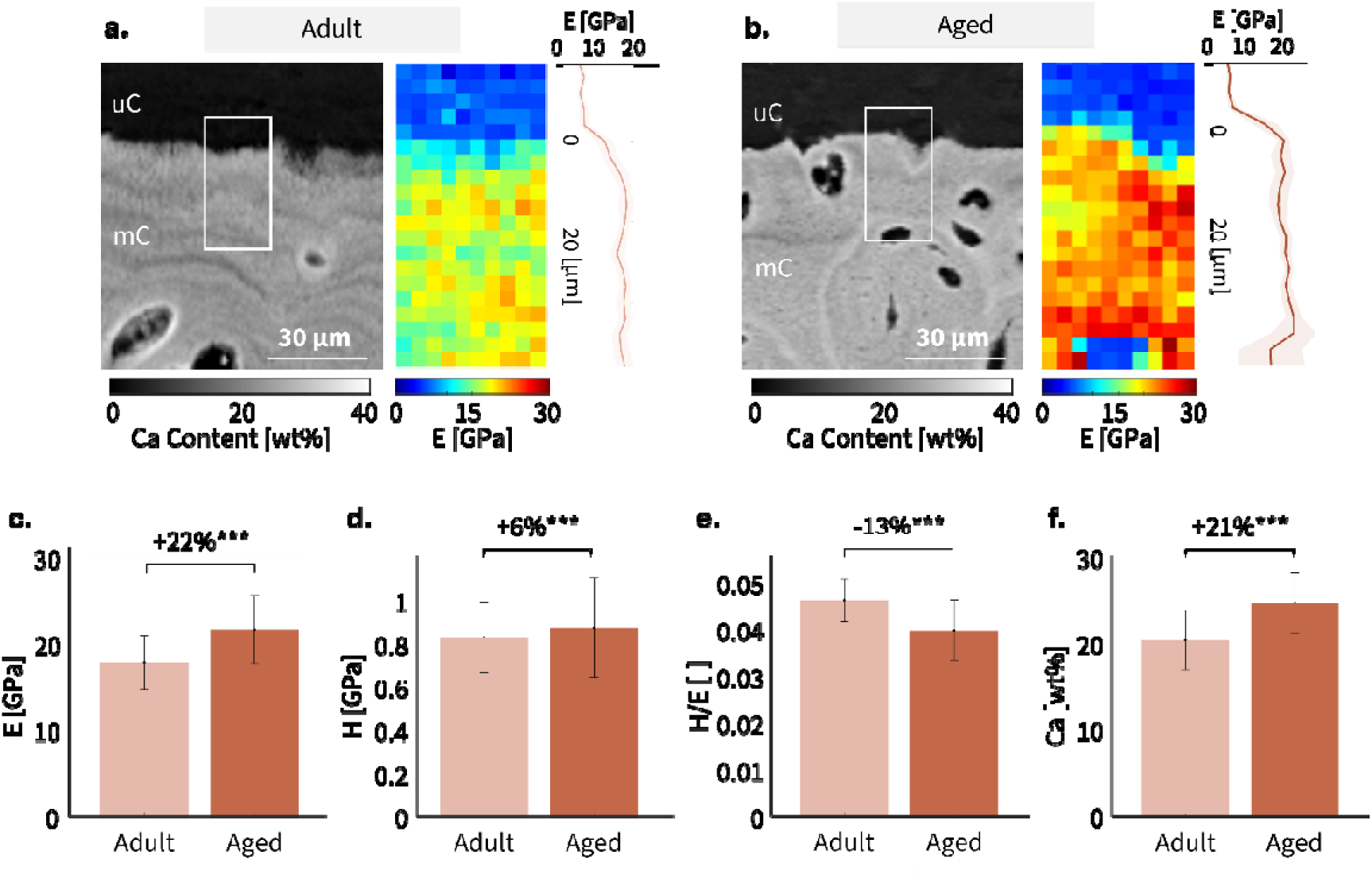
Mechanical characterization. Spatial maps of indentation modulus across the interface between unmineralized (uC) and mineralized cartilage (mC), and location-matched qBEI images showing the mineral content, for a representative (a) adult and (b) aged rats. The white frames highlight the indented area. The profile plots on the right side, computed on the 2D mechanical maps, underline the mechanical gradients at the interface. Quantitative comparison of (c) indentation modulus E, (d) hardness H, and (e) hardness/modulus ratio for adult and aged rats. (f) The corresponding region-matched mineral content is reported for interpreting the mechanical data. Data shown as mean (histogram height) with error bars representing one standard deviation. Statistical significance between data sets is indicated as follows: ***p < 0.001.

Other features present in the spatial patterns of mineral content were also reflected in the mechanical property maps. Former tidemarks, for example, are evident in qBEI maps as darker bands (hence having lower calcium), causing a drop in indentation modulus (Fig. 6a). The osteochondral cement line, appearing as a bright line in qBEI, had higher stiffness than neighboring tissue. Quantification of the mean indentation modulus and hardness in adult and aged samples revealed differences. Although both mechanical parameters increased with aging, the rise in modulus (+22 %, p < 0.001, Fig. 6c) was approximately three times greater than the increase in hardness (+6 %, p < 0.001, Fig. 6d). In the same regions, mineral content also increased by 21 % (p < 0.001, Fig. 6f). Consequently, the ratio between hardness and indentation modulus decreased significantly in aged animals (−13 %, p < 0.001, Fig. 6e). Similar trends were observed in subchondral bone (Supplementary Information, Fig. S8).

## Discussion

In this work, we combine multiple high-resolution techniques to investigate microstructural and material aspects at the osteochondral junction in aging rats. We specifically focus on mineralized cartilage and subchondral bone, with the former being so far under investigated. We found that these two tissues, despite being close neighbors, show specific microstructural and material alterations with aging. Starting from microstructural changes, we characterize subchondral bone at the epiphysis and metaphyseal bone away from the joint, distinguishing between cortical and trabecular compartments. In aged samples, both the subchondral plate (at the epiphysis) and the cortical bone (at the metaphysis) got thicker, with the former showing larger modifications. Recalling that aged rats were almost 50% heavier than adult ones (indicating continuous growth), the corresponding tibiae are likely exposed to higher loads. To withstand forces without too much deflection, a long bone needs, above all, high bending rigidity. At the metaphysis, this can probably be achieved by increasing the thickness of the cortical shell, by redistributing bone material farther from the neutral axis, or through a combination of both strategies [47]. In response to increased loading, the subchondral plate can limit deformation mainly by thickening, while placing material away from the neutral axis is not an option. Such constraints in mechanical adaptation, combined with different loading conditions and possibly other biological factors, may explain the more pronounced thickening of the subchondral plate compared to cortical bone. Plate thickening occurs mainly toward the underlying trabecular bone, most likely because bone formation is much easier there, due to the proximity to bone marrow and trabecular surfaces. The age-related thickening of the subchondral plate observed here agrees with previous studies on rodents [26,48]. In human individuals, subchondral bone has been investigated mainly in the context of OA. Thickening of this region happens also without weight gain, and it is considered a clear hallmark of this disease [49], yet whether it precedes or follows cartilage degradation is still not fully understood [50,51]. Concomitant with the increase in body weight, in aged rats we observed trabecular bone loss and coarsening of the trabecular network, the latter being somewhat less evident in the subchondral region. Age-related changes in subchondral bone may indeed be different compared to other skeletal locations due to the continuous mechanical loading from the joint. A previous investigation on subchondral bone microstructure in rats reported a continuous increase in trabecular BV/TV from 1 to 17 months of age [26], while at other locations trabecular bone may already undergo a gradual decline [52].

Turning to material aspects, the mineral content of subchondral bone and mineralized cartilage was heterogeneous and characterized by peak-shaped mineralization frequency distributions. In bone, the distribution of mineral content results from the interplay between mineralization and remodeling [29]: its shape and temporal evolution can therefore provide insight into these processes [53]. The finding that the mineralization frequency distribution in aged rats shifts towards higher mineral content suggests that the mineralization process prevails over remodeling, which is believed to be minimal in rats [46,54]. In subchondral bone, the rightward shift is accompanied by a higher peak and a somewhat narrower distribution, probably caused by a slowing down of the mineralization kinetics as the mineral content increases, leading to a “piling-up” effect [62]. Interestingly, such behavior was not observed in mineralized cartilage despite a higher increase in mineral content. This may be due to a much slower turnover of mineralized cartilage compared to subchondral bone. Essentially, old mineralized cartilage is occasionally replaced by subchondral bone close to the osteochondral cement line [30] and new mineralized cartilage is formed at the tidemark, where articular cartilage gets mineralized very slowly and in a discontinuous manner, as demonstrated by the presence of former tidemarks [27]. The absence of a clear “piling-up” effect may be due to the lower mineral content of this tissue being further away from the mineral content of fully mineralized cartilage (which may be as high as 30 wt % as inferred from the cartilaginous inclusions) or may even reflect a different mineralization velocity. Indeed, differences in the mineralization kinetics between the two tissues may potentially arise from intrinsic differences in the mineralization process (also considering the underlying organic matrix of collagen Type I in bone and Type II in mineralized cartilage) as well as from different ion transport pathways [55]. Active players of bone mineralization are osteocytes and their canaliculi, hosted in the so-called lacunocanalicular network (LCN) and contributing to the transport of mineral ions, precursors, and inhibitors of mineralization [41]. Cartilage mineralization relies on the active role of chondrocytes [56], which, however, do not form such a highly interconnected network but tend to aggregate into clusters [31], possibly to enable cell-cell interaction. The limited transport possibilities within mineralized cartilage may result in an overall slower mineralization, thereby explaining why mineralized cartilage has a lower mineral content than bone. However, if given sufficient time, the mineral content of this tissue can still increase and even exceed that of bone, as evidenced by the highly mineralized cartilaginous islands present in subchondral bone. In larger mammals (including humans), which have a longer lifespan, mineralized cartilage usually has a higher mineral content than bone [20,22,57]. One additional factor that may impact mineralization is the presence in both tissues of a nanoscale channel network, much smaller than the LCN, which may be further involved in the transport of ions and small molecules [58].

Examining mineral content heterogeneities, we found that regions around chondrocyte lacunae are not uniformly mineralized but present a thin hyper-mineralized layer close to the lacunae, more frequently seen in aged samples. Previous works have also reported regions of higher mineralization in the vicinity of chondrocyte lacunae [30,55], with the additional finding that the largest lacunae/clusters tend to be surrounded by more mineralized tissue [31]. The high degree of mineralization around the lacunae may be associated with a distinct organization of the perilacunar matrix in the form of concentric lamellae as observed in their immediate vicinity [59]. In aged rats, we reported a decrease in the size of chondrocyte lacunae, along with mineralized deposits in several lacunae. The decrease in size agrees with previous literature and may indicate a decline in cellular activity [60]. In bone, the occlusion of lacunae with minerals, named micropetrosis, is a well-documented phenomenon related to osteocyte death [30,61,62]. Conversely, mineral filling of chondrocyte lacunae within mineralized cartilage is still poorly understood, especially because this tissue is not transient as growth plate cartilage.

A central observation of our work is that aging modifies the gradient in mineral content present at the interface between unmineralized and mineralized cartilage. Not only did the width of the transition regions decrease by more than 50% in aged samples, but a higher mineralized band also appeared close to the mineralization front, possibly indicating mineral accumulation. In aged samples, markedly fewer arrested chondrocytes are observed at the unmineralized–mineralized cartilage interface; as these cells are thought to regulate mineralization at the tidemark, they may also contribute to maintaining a smooth transition zone. Performing nanoindentation at the same locations previously assessed with qBEI, we found that the gradient in mineral content across the tidemark is well reflected in a gradient in local elastic modulus, being steeper in aged samples. The local mechanical properties across the mineralization front have been analyzed in several previous works, and different transition widths, ranging from only a few micrometers to tens of micrometers, have been reported, depending on the anatomical location and experimental approach [9,12,20,22,23]. Of particular interest is that the higher mineral content in aged rats is associated with a large increase in elastic modulus but a much smaller increase in hardness. Assuming hardness as a rough indicator of yield stress [63], the ratio between hardness and elastic modulus can be informative on the tissue yield strain [64]. Our results show a decreased ratio in aged samples, suggesting a more brittle behavior. The presence of more cracks (probably due to sample preparation) in aged rats is an indirect confirmation of increased material-level fragility. While it is already known that yield strain decreases in aged bone [65], one contributing factor being non-enzymatic collagen cross-links [66], here we complement such knowledge by showing that mineralized cartilage also becomes weaker with aging.

The final characterization step of our multimodal approach involved SHG imaging, which underlined striking differences in the collagen fibril arrangement between mineralized cartilage and bone. In agreement with previous observations [67–69], the absence of a clear lamellar pattern, combined with a more uniform texture, should indicate that in mineralized cartilage, collagen fibrils are not assembled into higher-order lamellar motifs and are much more uniformly arranged along a common direction than in bone. It has also been suggested that collagen Type II may form finer fibrils than Type I, reflected into different SHG signatures [67,68]. The generally very low SHG signal at the osteochondral cement line may indicate that fibril organization at this interface is quite different from neighboring tissues. This is in line with previous works focusing on the cement line around osteons, analyzed with SHG and X-ray scattering [70,71] as well as with focused ion beam - scanning electron microscopy (FIB-SEM) [72]. One additional interesting finding is that, when visible, subchondral bone lamellae adjacent to the osteochondral cement line tend to be oriented parallel to it. A similar pattern was found at other anatomical locations [24], even including the bone-tendon junction [12]. This further underlines the role of the cement line as a rigid substrate for the coordinate deposition of lamellar bone [73]. As the cement line seems to colocalize with discontinuous fibril arrangement, our data further support the idea that anchoring between mineralized cartilage and bone strongly relies on interlocking between the two tissues. Nonetheless, some fibrils crossing the osteochondral cement line have also been observed at a different anatomical location (e.g., metacarpophalangeal joint) [67].

Concerning limitations, it should be noted that the number of tested specimens per group varied depending on the characterization method (10 for micro-CT, 6 for qBEI, and 3 for nIND/SHG). While sample preparation for micro-CT is fast and uncomplicated, the other techniques require more time-consuming sample preparation and (for nIND) measurement. It is common that, when moving from lower- to higher-resolution methods, the field of view decreases. Given the high heterogeneity of material-level properties [9,12,18,31,46], laboratory rodents usually show greater intra-individual than inter-individual variability [74]. Therefore, we decided to characterize material aspects in fewer samples but in more detail. The resulting mechanical properties and fibril orientation were consistent across all specimens, and we could detect statistically significant differences in mechanical properties between different tissues and age groups, well reflecting changes in mineral content measured on more samples. We should also mention that samples were tested dehydrated, although we know that this has an impact on the mechanical properties. Dehydration with ethanol is believed to remove water while maintaining (if not increasing) hydrogen-hydrogen bonding in collagen [75]. Removing water typically leads to an increase in indentation modulus and hardness [76]. As our focus is primarily on relative differences between mineralized cartilage and bone and between adult and aged conditions, we would expect dehydration to affect these aspects in a comparable manner, also considering that samples were prepared under identical conditions. Two advantages of dehydration include the possibility to perform correlative mechanics-mineral content measurements on the same samples and locations, as well as to avoid reproducibility issues, as in hydrated samples, mechanical testing becomes very sensitive to variations of hydration state. [77,78]. These findings from a preclinical model provide new insights into the osteochondral junction, which remain to be confirmed in humans.

## Conclusions

Characterizing physiological changes during aging at the osteochondral junction is very critical to understand how aging can contribute to degenerative joint disease. Using a multimodal correlative approach allowing to evaluate alterations in microstructure, mineral content, collagen organization, and local mechanical properties, we highlight that mineralized cartilage and subchondral bone are affected to different degrees by the aging process. Microstructure and material level changes both suggest an overall increase in the rigidity of the mineralized tissues underlying articular cartilage. This is a critical aspect as these alterations may limit the deformation and the energy absorption capacity of the epiphysis, thereby placing additional burden on articular cartilage and increasing the probability of tissue degeneration.

## Acknowledgements

L.M. obtained funding from F.R.S.-FNRS as part of a FRIA grant (n. FC 47333). L.M. H.v.L. and D.R. are grateful for financial support from the Excellence of Science (EOS) program (EOS no. 40007553). We warmly thank Pierre Drion and Luc Duwez from the University of Liege, GIGA, for the essential help to obtain the samples and for sample extraction. We wish to thank Fahimeh Azari and Jilmen Quintiens for taking the micro-CT scans at FIBEr, KU Leuven. We thank Petra Keplinger, Sonja Lueger, and Phaedra Messmer from LBIO for the great help with image acquisition. MH and SB are grateful for financial support from the AUVA (Research funds of the Austrian workers’ compensation board) and OEGK (Austrian Social Health Insurance Fund). RW acknowledges support from the Max Planck Queensland Centre for the Materials Science of Extracellular Matrices.

## References

[1] C.D. Hoemann, C.-H. Lafantaisie-Favreau, V. Lascau-Coman, G. Chen, J. Guzmán-Morales, The cartilage-bone interface, J. Knee Surg. 25 (2012) 85–97. 10.1055/s-0032-1319782.

[2] X. Fan, X. Wu, R. Crawford, Y. Xiao, I. Prasadam, Macro, Micro, and Molecular. Changes of the Osteochondral Interface in Osteoarthritis Development, Front. Cell Dev. Biol. 9 (2021). 10.3389/fcell.2021.659654.

[3] S.I.M. Lepage, N. Robson, H. Gilmore, O. Davis, A. Hooper, S. St. John, V. Kamesan, P. Gelis, D. Carvajal, M. Hurtig, T.G. Koch, Beyond Cartilage Repair: The Role of the Osteochondral Unit in Joint Health and Disease, Tissue Eng. Part B Rev. 25 (2019) 114–125. 10.1089/ten.teb.2018.0122.

[4] L. Zhou, V.O. Gjvm, J. Malda, M.J. Stoddart, Y. Lai, R.G. Richards, K. Ki-Wai Ho, L. Qin, Innovative Tissue-Engineered Strategies for Osteochondral Defect Repair and Regeneration: Current Progress and Challenges, Adv. Healthc. Mater. 9 (2020) e2001008. 10.1002/adhm.202001008.

[5] J.W.C. Dunlop, R. Weinkamer, P. Fratzl, Artful interfaces within biological materials, Mater. Today 14 (2011) 70–78. 10.1016/S1369-7021(11)70056-6.

[6] S. Thomopoulos, V. Birman, G.M. Genin, Structural Interfaces and Attachments in Biology, Springer Science & Business Media, 2012.

[7] Z. Su, P. Tan, J. Zhang, P. Wang, S. Zhu, N. Jiang, Understanding the Mechanics of the Temporomandibular Joint Osteochondral Interface from Micro-and Nanoscopic Perspectives, Nano Lett. 23 (2023) 11702–11709. 10.1021/acs.nanolett.3c03564.

[8] W. Badar, S.R. Inamdar, P. Fratzl, T. Snow, N.J. Terrill, M.M. Knight, H.S. Gupta, Nonlinear Stress-Induced Transformations in Collagen Fibrillar Organization, Disorder and Strain Mechanisms in the Bone-Cartilage Unit, Adv. Sci. 12 (2025) 2407649. 10.1002/advs.202407649.

[9] S.E. Campbell, V.L. Ferguson, D.C. Hurley, Nanomechanical mapping of the osteochondral interface with contact resonance force microscopy and nanoindentation, Acta Biomater. 8 (2012) 4389–4396. 10.1016/j.actbio.2012.07.042.

[10] X. Wang, J. Lin, Z. Li, Y. Ma, X. Zhang, Q. He, Q. Wu, Y. Yan, W. Wei, X. Yao, C. Li, W. Li, S. Xie, Y. Hu, S. Zhang, Y. Hong, X. Li, W. Chen, W. Duan, H. Ouyang, Identification of an Ultrathin Osteochondral Interface Tissue with Specific Nanostructure at the Human Knee Joint, Nano Lett. 22 (2022) 2309–2319. 10.1021/acs.nanolett.1c04649.

[11] A. Moayedi, K. Karali, M. Boese, J. Radulovic, G. Blunn, 3D full-field lacunar morphology and deformation of calcified fibrocartilage in the loaded Achilles enthesis of a mouse, Commun. Mater. 6 (2025) 249. 10.1038/s43246-025-00972-3.

[12] A. Tits, S. Blouin, M. Rummler, J.-F. Kaux, P. Drion, G.H. van Lenthe, R. Weinkamer, M.A. Hartmann, D. Ruffoni, Structural and functional heterogeneity of mineralized fibrocartilage at the Achilles tendon-bone insertion, Acta Biomater. 166 (2023) 409–418. 10.1016/j.actbio.2023.04.018.

[13] A. Tits, E. Plougonven, S. Blouin, M.A. Hartmann, J.-F. Kaux, P. Drion, J. Fernandez, G.H. van Lenthe, D. Ruffoni, Local anisotropy in mineralized fibrocartilage and subchondral bone beneath the tendon-bone interface, Sci. Rep. 11 (2021) 16534. 10.1038/s41598-021-95917-4.

[14] A. Tits, D. Ruffoni, Joining soft tissues to bone: Insights from modeling and simulations, Bone Rep. 14 (2021) 100742. 10.1016/j.bonr.2020.100742.

[15] L.A.E. Evans, A.A. Pitsillides, Structural clues to articular calcified cartilage function: A descriptive review of this crucial interface tissue, J. Anat. 241 (2022) 875–895. 10.1111/joa.13728.

[16] K. Madi, K.A. Staines, B.K. Bay, B. Javaheri, H. Geng, A.J. Bodey, S. Cartmell, A.A. Pitsillides, P.D. Lee, In situ characterization of nanoscale strains in loaded whole joints via synchrotron X-ray tomography, Nat. Biomed. Eng. 4 (2020) 343–354. 10.1038/s41551-019-0477-1.

[17] J. Pan, X. Zhou, W. Li, J.E. Novotny, S.B. Doty, L. Wang, In situ measurement of transport between subchondral bone and articular cartilage, J. Orthop. Res. 27 (2009) 1347–1352. 10.1002/jor.20883.

[18] I. Zizak, P. Roschger, O. Paris, B.M. Misof, A. Berzlanovich, S. Bernstorff, H. Amenitsch, K. Klaushofer, P. Fratzl, Characteristics of mineral particles in the human bone/cartilage interface, J. Struct. Biol. 141 (2003) 208–217. 10.1016/s1047-8477(02)00635-4.

[19] M.A.J. Finnilä, S. Das Gupta, M.J. Turunen, I. Hellberg, A. Turkiewicz, V. Lutz_-_Bueno, E. Jonsson, M. Holler, N. Ali, V. Hughes, H. Isaksson, J. Tjörnstrand, P. Önnerfjord, M. Guizar_-_Sicairos, S. Saarakkala, M. Englund, Mineral Crystal Thickness in Calcified Cartilage and Subchondral Bone in Healthy and Osteoarthritic Human Knees, J. Bone Miner. Res. 37 (2022) 1700–1710. 10.1002/jbmr.4642.

[20] H.S. Gupta, S. Schratter, W. Tesch, P. Roschger, A. Berzlanovich, T. Schoeberl, K. Klaushofer, P. Fratzl, Two different correlations between nanoindentation modulus and mineral content in the bone-cartilage interface, J. Struct. Biol. 149 (2005) 138–148. 10.1016/j.jsb.2004.10.010.

[21] F. Wang, Z. Ying, X. Duan, H. Tan, B. Yang, L. Guo, G. Chen, G. Dai, Z. Ma, L. Yang, Histomorphometric analysis of adult articular calcified cartilage zone, J. Struct. Biol. 168 (2009) 359–365. 10.1016/j.jsb.2009.08.010.

[22] V.L. Ferguson, A.J. Bushby, A. Boyde, Nanomechanical properties and mineral concentration in articular calcified cartilage and subchondral bone, J. Anat. 203 (2003) 191–202. 10.1046/j.1469-7580.2003.00193.x.

[23] M. Doube, E.C. Firth, A. Boyde, A.J. Bushby, Combined nanoindentation testing and scanning electron microscopy of bone and articular calcified cartilage in an equine fracture predilection site, Eur. Cell. Mater. 19 (2010) 242–251. 10.22203/ecm.v019a23.

[24] J. Hu, K. Zheng, B.E. Sherlock, J. Zhong, J. Mansfield, E. Green, A.D. Toms, C.P. Winlove, J. Chen, Zonal Characteristics of Collagen Ultrastructure and Responses to Mechanical Loading in Articular Cartilage, Acta Biomater. 195 (2025) 104–116. 10.1016/j.actbio.2025.01.047.

[25] K. Hayat, N. Marr, K.K.L. Mak, M. Doube, Type-I and -II collagens from bone and cartilage colocalize at the osteochondral cement line, Bone Jt. Res. 14 (2025) 735–744. 10.1302/2046-3758.148.BJR-2024-0396.R1.

[26] P. Ren, H. Niu, H. Gong, R. Zhang, Y. Fan, Morphological, biochemical and mechanical properties of articular cartilage and subchondral bone in rat tibial plateau are age related, J. Anat. 232 (2018) 457–471. 10.1111/joa.12756.

[27] M. Doube, E.C. Firth, A. Boyde, Variations in articular calcified cartilage by site and exercise in the 18-month-old equine distal metacarpal condyle, Osteoarthritis Cartilage 15 (2007) 1283–1292. 10.1016/j.joca.2007.04.003.

[28] P. Roschger, E.P. Paschalis, P. Fratzl, K. Klaushofer, Bone mineralization density distribution in health and disease, Bone 42 (2008) 456–466. 10.1016/j.bone.2007.10.021.

[29] D. Ruffoni, P. Fratzl, P. Roschger, K. Klaushofer, R. Weinkamer, The bone mineralization density distribution as a fingerprint of the mineralization process, Bone 40 (2007) 1308–1319. 10.1016/j.bone.2007.01.012.

[30] A. Boyde, The Bone Cartilage Interface and Osteoarthritis, Calcif. Tissue Int. 109 (2021) 303–328. 10.1007/s00223-021-00866-9.

[31] T. Tang, J. Zhong, J. Hu, V. Schemenz, A. Davydok, R. Brunner, J. Zhou, W. Wagermaier, A.A. Pitsillides, W.J. Landis, P. Fratzl, J. Chen, Gradients in lacunar morphology and cartilage mineralization reflect the mechanical function of the mouse femoral head epiphysis, Acta Biomater. 201 (2025) 385–399. 10.1016/j.actbio.2025.06.002.

[32] N. Otsu, A Threshold Selection Method from Gray-Level Histograms, IEEE Trans. Syst. Man Cybern. 9 (1979) 62–66. 10.1109/TSMC.1979.4310076.

[33] M.L. Bouxsein, S.K. Boyd, B.A. Christiansen, R.E. Guldberg, K.J. Jepsen, R. Müller, Guidelines for assessment of bone microstructure in rodents using micro-computed tomography, J. Bone Miner. Res. 25 (2010) 1468–1486. 10.1002/jbmr.141.

34. J. Maier, M. Black, S. Bonaretti, B. Bier, B. Eskofier, J.-H. Choi, M. Levenston, G. Gold, R. Fahrig, A. Maier, Comparison of Different Approaches for Measuring Tibial Cartilage Thickness, J. Integr. Bioinforma. 14 (2017) 20170015. 10.1515/jib-2017-0015.

[35] B. Schamberger, R. Ziege, K. Anselme, M. Ben Amar, M. Bykowski, A.P.G. Castro, A. Cipitria, R.A. Coles, R. Dimova, M. Eder, S. Ehrig, L.M. Escudero, M.E. Evans, P.R. Fernandes, P. Fratzl, L. Geris, N. Gierlinger, E. Hannezo, A. Iglič, J.J.K. Kirkensgaard, P. Kollmannsberger, Ł. Kowalewska, N.A. Kurniawan, I. Papantoniou, L. Pieuchot, T.H.V. Pires, L.D. Renner, A.O. Sageman-Furnas, G.E. Schröder-Turk, A. Sengupta, V.R. Sharma, A. Tagua, C. Tomba, X. Trepat, S.L. Waters, E.F. Yeo, A. Roschger, C.M. Bidan, J.W.C. Dunlop, Curvature in Biological Systems: Its Quantification, Emergence, and Implications across the Scales, Adv. Mater. Deerfield Beach Fla 35 (2023) e2206110. 10.1002/adma.202206110.

[36] T. Hildebrand, P. Rüegsegger, A new method for the model-independent assessment of thickness in three-dimensional images, J. Microsc. 185 (1997) 67–75. 10.1046/j.1365-2818.1997.1340694.x.

[37] P. Roschger, P. Fratzl, J. Eschberger, K. Klaushofer, Validation of quantitative backscattered electron imaging for the measurement of mineral density distribution in human bone biopsies, Bone 23 (1998) 319–326. 10.1016/s8756-3282(98)00112-4.

[38] P. Roschger, J. Eschberger, H. Plenk, Formation of Ultracracks in Methacrylate-Embedded Undecalcified Bone Samples by Exposure to Aqueous Solutions, Cells Mater. 3 (1993). https://digitalcommons.usu.edu/cellsandmaterials/vol3/iss4/3.

[39] M.A. Hartmann, S. Blouin, B.M. Misof, N. Fratzl-Zelman, P. Roschger, A. Berzlanovich, G.M. Gruber, P.C. Brugger, J. Zwerina, P. Fratzl, Quantitative Backscattered Electron Imaging of Bone Using a Thermionic or a Field Emission Electron Source, Calcif. Tissue Int. 109 (2021) 190–202. 10.1007/s00223-021-00832-5.

[40] C. Lukas, P. Kollmannsberger, D. Ruffoni, P. Roschger, P. Fratzl, R. Weinkamer, The Heterogeneous Mineral Content of Bone—Using Stochastic Arguments and Simulations to Overcome Experimental Limitations, J. Stat. Phys. 144 (2011) 316–331. 10.1007/s10955-011-0209-8.

[41] M. Ayoubi, A.F. van Tol, R. Weinkamer, P. Roschger, P.C. Brugger, A. Berzlanovich, L. Bertinetti, A. Roschger, P. Fratzl, 3D Interrelationship between Osteocyte Network and Forming Mineral during Human Bone Remodeling, Adv. Healthc. Mater. 10 (2021) 2100113. 10.1002/adhm.202100113.

[42] Q. Kan, W. Yan, G. Kang, Q. Sun, Oliver–Pharr indentation method in determining elastic moduli of shape memory alloys—A phase transformable material, J. Mech. Phys. Solids 61 (2013) 2015–2033. 10.1016/j.jmps.2013.05.007.

[43] R.M. Williams, W.R. Zipfel, W.W. Webb, Interpreting Second-Harmonic Generation Images of Collagen I Fibrils, Biophys. J. 88 (2005) 1377–1386. 10.1529/biophysj.104.047308.

[44] X. Chen, O. Nadiarynkh, S. Plotnikov, P.J. Campagnola, Second harmonic generation microscopy for quantitative analysis of collagen fibrillar structure, Nat. Protoc. 7 (2012) 654–669. 10.1038/nprot.2012.009.

[45] M.-A. Houle, C.-A. Couture, S. Bancelin, J. Van der Kolk, E. Auger, C. Brown, K. Popov, L. Ramunno, F. Légaré, Analysis of forward and backward Second Harmonic Generation images to probe the nanoscale structure of collagen within bone and cartilage, J. Biophotonics 8 (2015) 993–1001. 10.1002/jbio.201500150.

[46] A. Shipov, P. Zaslansky, H. Riesemeier, G. Segev, A. Atkins, R. Shahar, Unremodeled endochondral bone is a major architectural component of the cortical bone of the rat (*Rattus norvegicus*), J. Struct. Biol. 183 (2013) 132–140. 10.1016/j.jsb.2013.04.010.

[47] J. Jast, I. Jasiuk, Age-related changes in the 3D hierarchical structure of rat tibia cortical bone characterized by high-resolution micro-CT., J. Appl. Physiol. Bethesda Md 1985 114 (2013) 923–933. 10.1152/japplphysiol.00948.2011.

[48] N. Hamann, G.-P. Brüggemann, A. Niehoff, Topographical variations in articular cartilage and subchondral bone of the normal rat knee are age-related, Ann. Anat. Anat. Anz. Off. Organ Anat. Ges. 196 (2014) 278–285. 10.1016/j.aanat.2014.04.006.

[49] B. Gatenholm, C. Lindahl, M. Brittberg, V.A. Stadelmann, Spatially matching morphometric assessment of cartilage and subchondral bone in osteoarthritic human knee joint with micro-computed tomography, Bone 120 (2019) 393–402. 10.1016/j.bone.2018.12.003.

[50] D.B. Burr, The importance of subchondral bone in osteoarthrosis, Curr. Opin. Rheumatol. 10 (1998) 256.

[51] A.J. Bailey, J.P. Mansell, Do Subchondral Bone Changes Exacerbate or Precede Articular Cartilage Destruction in Osteoarthritis of the Elderly?, Gerontology 43 (2009) 296–304. 10.1159/000213866.

[52] R. Zhang, H. Gong, D. Zhu, R. Ma, J. Fang, Y. Fan, Multi-level femoral morphology and mechanical properties of rats of different ages, Bone 76 (2015) 76–87. 10.1016/j.bone.2015.03.022.

[53] D. Ruffoni, P. Fratzl, P. Roschger, R. Phipps, K. Klaushofer, R. Weinkamer, Effect of Temporal Changes in Bone Turnover on the Bone Mineralization Density Distribution: A Computer Simulation Study, J. Bone Miner. Res. 23 (2008) 1905–1914. 10.1359/jbmr.080711.

[54] R.G. Erben, Trabecular and endocortical bone surfaces in the rat: Modeling or remodeling?, Anat. Rec. 246 (1996) 39–46. 10.1002/(SICI)1097-0185(199609)246:1%253C39::AID-AR5%253E3.0.CO;2-A.

[55] P.A. Zecca, M. Reguzzoni, M. Protasoni, M. Raspanti, The chondro-osseous junction of articular cartilage, Tissue Cell 80 (2023) 101993. 10.1016/j.tice.2022.101993.

[56] S. Boonrungsiman, C. Allen, F. Nudelman, S. Shefelbine, C. Farquharson, A.E. Porter, R.A. Fleck, Endochondral ossification: Insights into the cartilage mineralization processes achieved by an anhydrous freeze substitution protocol, Acta Biomater. 191 (2025) 149–157. 10.1016/j.actbio.2024.11.015.

[57] A. Sensini, L. Raimondi, A. Malerba, C.P. Silva, A. Zucchelli, A. Tits, D. Ruffoni, S. Blouin, M.A. Hartmann, M. van Griensven, L. Moroni, Understanding the structure and mechanics of the sheep calcaneal enthesis: a relevant animal model to design scaffolds for tissue engineering applications, Biomater. Adv. 175 (2025) 214320. 10.1016/j.bioadv.2025.214320.

[58] T. Tang, W. Landis, E. Raguin, P. Werner, L. Bertinetti, M. Dean, W. Wagermaier, P. Fratzl, A 3D Network of Nanochannels for Possible Ion and Molecule Transit in Mineralizing Bone and Cartilage, Adv. NanoBiomed Res. 2 (2022) 2100162. 10.1002/anbr.202100162.

[59] C.M. Keenan, A.J. Beckett, H. Sutherland, L.R. Ranganath, J.C. Jarvis, I.A. Prior, J.A. Gallagher, Concentric lamellae – novel microanatomical structures in the articular calcified cartilage of mice, Sci. Rep. 9 (2019) 11188. 10.1038/s41598-019-47545-2.

[60] J.Y. Oda, E.A. Liberti, L.B.M. Maifrino, R.R. de Souza, Variation in articular cartilage in rats between 3 and 32 months old. A histomorphometric and scanning electron microscopy study, Biogerontology 8 (2007) 345–352. 10.1007/s10522-006-9076-0.

61. [61] P. Milovanovic, B. Busse, Phenomenon of osteocyte lacunar mineralization: indicator of former osteocyte death and a novel marker of impaired bone quality?, (2020). 10.1530/EC-19-0531.

[62] S. Blouin, B.M. Misof, M. Mähr, N. Fratzl-Zelman, P. Roschger, S. Lueger, P. Messmer, P. Keplinger, F. Rauch, F.H. Glorieux, A. Berzlanovich, G.M. Gruber, P.C. Brugger, E. Shane, R.R. Recker, J. Zwerina, M.A. Hartmann, Osteocyte lacunae in transiliac bone biopsy samples across life span, Acta Biomater. 157 (2023) 275–287. 10.1016/j.actbio.2022.11.051.

[63] A. Ibrahim, N. Magliulo, J. Groben, A. Padilla, F. Akbik, Z.A. Hamid, Hardness, an Important Indicator of Bone Quality, and the Role of Collagen in Bone Hardness, J. Funct. Biomater. 11 (2020). 10.3390/jfb11040085.

[64] H. Rojacz, A. Nevosad, M. Varga, On wear mechanisms and microstructural changes in nano-scratches of fcc metals, Wear 526–527 (2023) 204928. 10.1016/j.wear.2023.204928.

[65] L. Ravazzano, G. Colaianni, A. Tarakanova, Y.-B. Xiao, M. Grano, F. Libonati, Multiscale and multidisciplinary analysis of aging processes in bone, Npj Aging 10 (2024) 28. 10.1038/s41514-024-00156-2.

[66] J.S. Nyman, A. Roy, J.H. Tyler, R.L. Acuna, H.J. Gayle, X. Wang, Age-related factors affecting the postyield energy dissipation of human cortical bone, J. Orthop. Res. 25 (2007) 646–655. 10.1002/jor.20337.

[67] J.C. Mansfield, C. Peter Winlove, A multi-modal multiphoton investigation of microstructure in the deep zone and calcified cartilage, J. Anat. 220 (2012) 405–416. 10.1111/j.1469-7580.2012.01479.x.

[68] T. Saitou, H. Kiyomatsu, T. Imamura, Quantitative Morphometry for Osteochondral Tissues Using Second Harmonic Generation Microscopy and Image Texture Information, Sci. Rep. 8 (2018) 2826. 10.1038/s41598-018-21005-9.

[69] J. Hu, K. Zheng, B.E. Sherlock, J. Zhong, J. Mansfield, E. Green, A.D. Toms, C.P. Winlove, J. Chen, Zonal Characteristics of Collagen Ultrastructure and Responses to Mechanical Loading in Articular Cartilage, Acta Biomater. 195 (2025) 104–116. 10.1016/j.actbio.2025.01.047.

[70] A. Cantamessa, V. Schemenz, S. Blouin, T. Volders, S. Amini, M. Rummler, J. Liu, A. Berzlanovich, R. Weinkamer, W. Wagermaier, M.A. Hartmann, D. Ruffoni, Cement Lines are Stiffer and Harder than Bone but Exhibit Different Mineral–Mechanics Relationships due to Thicker and Shorter Mineral Particles, Small Struct. (2025). 10.1002/sstr.202500612.

[71] A. Cantamessa, S. Blouin, M. Rummler, A. Berzlanovich, R. Weinkamer, M.A. Hartmann, D. Ruffoni, The mineralization of osteonal cement line depends on where the osteon is formed, JBMR Plus 9 (2025) ziaf114. 10.1093/jbmrpl/ziaf114.

[72] E. Raguin, K. Rechav, R. Shahar, S. Weiner, Focused ion beam-SEM 3D analysis of mineralized osteonal bone: lamellae and cement sheath structures, Acta Biomater. 121 (2021) 497–513. 10.1016/j.actbio.2020.11.002.

[73] M. Kerschnitzki, W. Wagermaier, P. Roschger, J. Seto, R. Shahar, G.N. Duda, S. Mundlos, P. Fratzl, The organization of the osteocyte network mirrors the extracellular matrix orientation in bone, J. Struct. Biol. 173 (2011) 303–311. 10.1016/j.jsb.2010.11.014.

[74] M. Rummler, A. van Tol, V. Schemenz, M.A. Hartmann, S. Blouin, B.M. Willie, R. Weinkamer, The Lacunocanalicular Network is Denser in C57BL/6 Compared to BALB/c Mice, Calcif. Tissue Int. 115 (2024) 744–758. 10.1007/s00223-024-01289-y.

[75] M. Granke, M.D. Does, J.S. Nyman, The Role of Water Compartments in the Material Properties of Cortical Bone, Calcif. Tissue Int. 97 (2015) 292–307. 10.1007/s00223-015-9977-5.

[76] A.J. Bushby, V.L. Ferguson, A. Boyde, Nanoindentation of bone: Comparison of specimens tested in liquid and embedded in polymethylmethacrylate, J. Mater. Res. 19 (2004) 249–259. 10.1557/jmr.2004.19.1.249.

[77] A. Faingold, S.R. Cohen, R. Shahar, S. Weiner, L. Rapoport, H.D. Wagner, The effect of hydration on mechanical anisotropy, topography and fibril organization of the osteonal lamellae, J. Biomech. 47 (2014) 367–372. 10.1016/j.jbiomech.2013.11.022.

[78] J.S. Nyman, A. Roy, X. Shen, R.L. Acuna, J.H. Tyler, X. Wang, The influence of water removal on the strength and toughness of cortical bone, J. Biomech. 39 (2006) 931–938. 10.1016/j.jbiomech.2005.01.012.

