## Supplementary material for "Aging modifies microstructure and material properties of mineralized cartilage and subchondral bone in the murine knee": SI

**Supplementary Information**

**Materials and Methods**

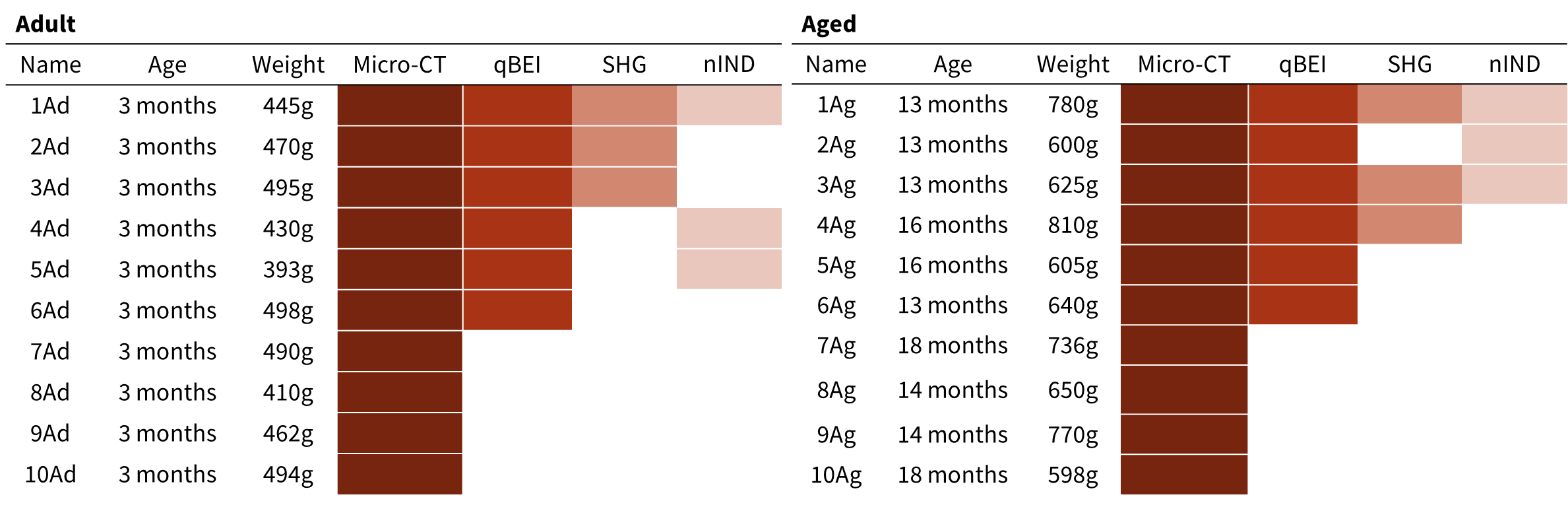

Table S1: Information on the Wistar rat samples used in this study, including animal age, weight, and the experimental technique employed for the characterization.

*Supplementary note 1: Tibia alignment and trabecular segmentation*

Tibia alignment was performed by registering all specimens to a common reference sample. One adult tibia was first aligned along the principal axes of inertia of its diaphysis using BoneJ [1] and cropped to retain a region extending from the tibial plateau to the onset of the medullary cavity, with a total length of 1 cm. All other tibiae were then rigidly registered to this reference sample using a mutual information metric and resampled with Lanczos interpolation (Avizo, Thermo Fisher Scientific). The segmentation of the cortical and trabecular compartments followed a semiautomatic workflow, as detailed in Fig. S1.

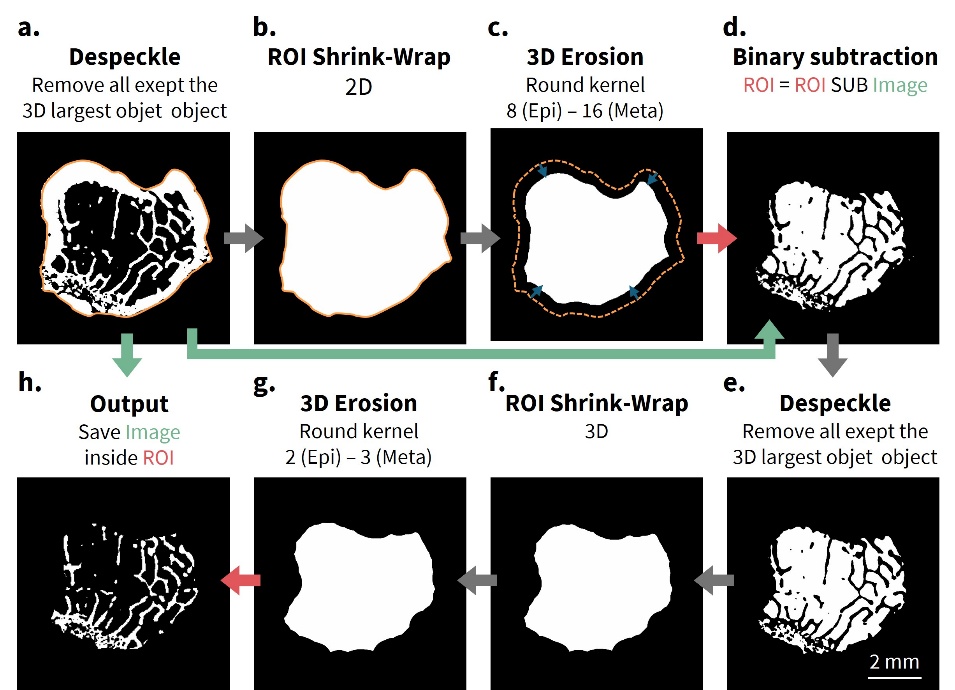

Figure S1: Semi-automated workflow for segmentation of trabecular and cortical compartments at the metaphysis (an analogous workflow was applied at the epiphysis). (a) Prior to segmentation, the virtual stacks were resliced to ensure that trabecular bone was always enclosed by a visible continuous cortical shell. A first despeckle operator was applied to remove isolated components. (b) The outer bone contour was then generated using a shrink-wrap operator and (c) eroded (8 and 16 pixels in the epiphysis and metaphysis, respectively). (d) A mask of the eroded contour was then subtracted from the “despeckled” bone volume to obtain a mask of the marrow space. (e) A despeckle operator was applied to the obtained mask. (f) The external contour of the marrow space was generated using the same shrink-wrap procedure and eroded (2 and 3 pixels in the epiphysis and metaphysis, respectively). (h) The resulting mask was used to extract the trabecular compartment.

*Supplementary note 2: Segmentation and smoothing of the subchondral plate*

To isolate the subchondral plate, masks were manually generated considering the projection of the epiphysis on the transverse plane to delineate the regions of interest in the medial and lateral plateaus. Within each mask, the subchondral plate located directly beneath the articular surface was extracted. Prior to computing plate thickness, the extracted volumes were smoothed to reduce roughness arising from small portions of trabecular bone attached to the cortical plate and not removed during masking. Smoothing was carried out in CTAn by applying four iterations of a 3D Gaussian filter (round kernel, size 3), followed by Otsu’s automatic binarization.

**Results**

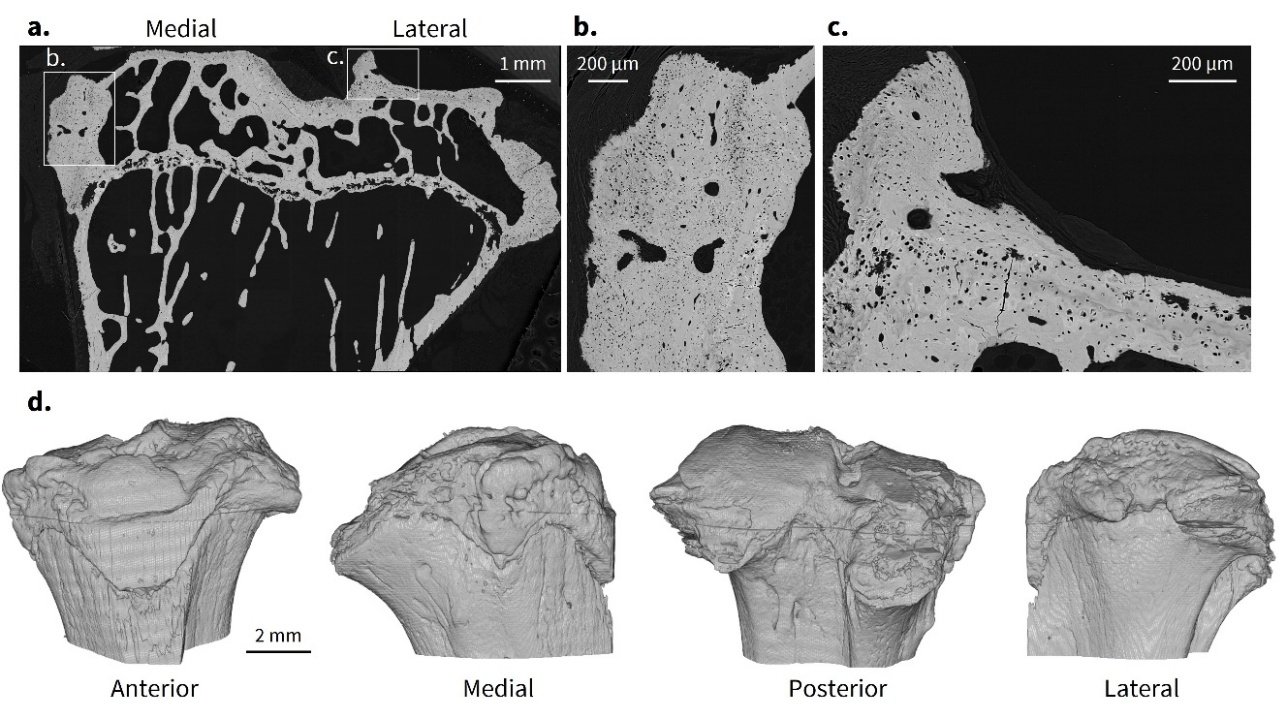

Figure S2: Sample with signs of joint degeneration. One tibia from the 15-month-old group was excluded from the analyses as it shows both microstructural and material alterations. (a) qBEI image of a mid-frontal section shows sparse trabecular bone with thin, poorly connected, and highly spaced trabeculae. Marked cortical thickening is evident on both medial and lateral sides of the epiphysis. (b) Magnified view of cortical thickening at the medial side. (c) Magnified view highlighting a layer of bone overlying mineralized cartilage, probably terminating into a bone spur. (d) Micro-CT scan reveals a striking shape alteration of the medial plateau.

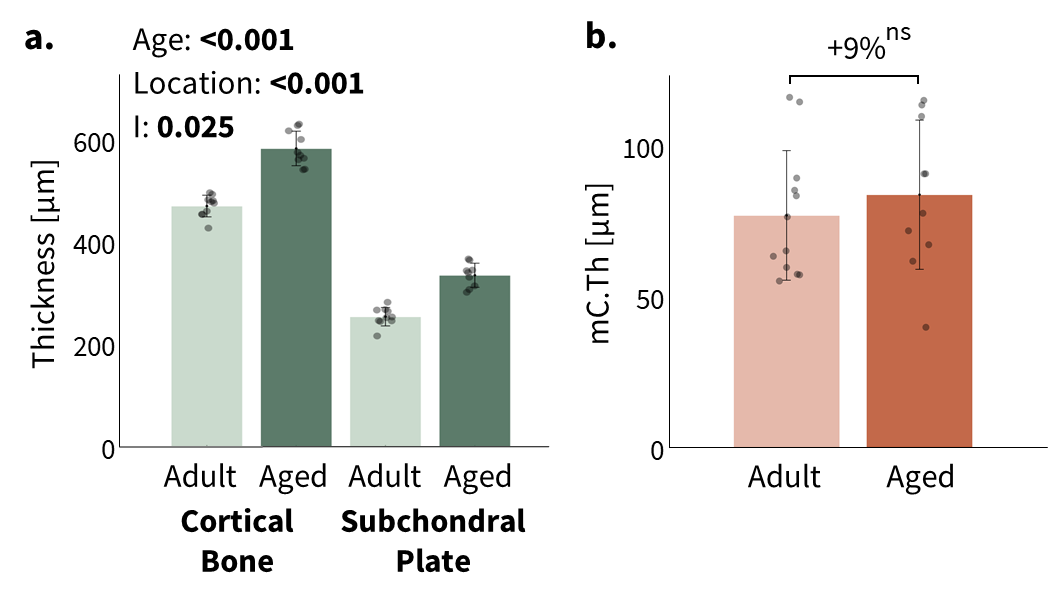

Figure S3: (a) Comparison of cortical thickness between adult and aged rats at the metaphyseal cortical bone versus the subchondral bone plate. (b) Thickness of mineralized cartilage in adult and aged rats. Data shown as mean (histogram height) with bars representing one standard deviation. Individual samples are reported as scattered points. For bone, results of two-way ANOVA are reported considering age, location and their interaction (I). For mineralized cartilage, results of the t-test are reported with ns indicating not significant.

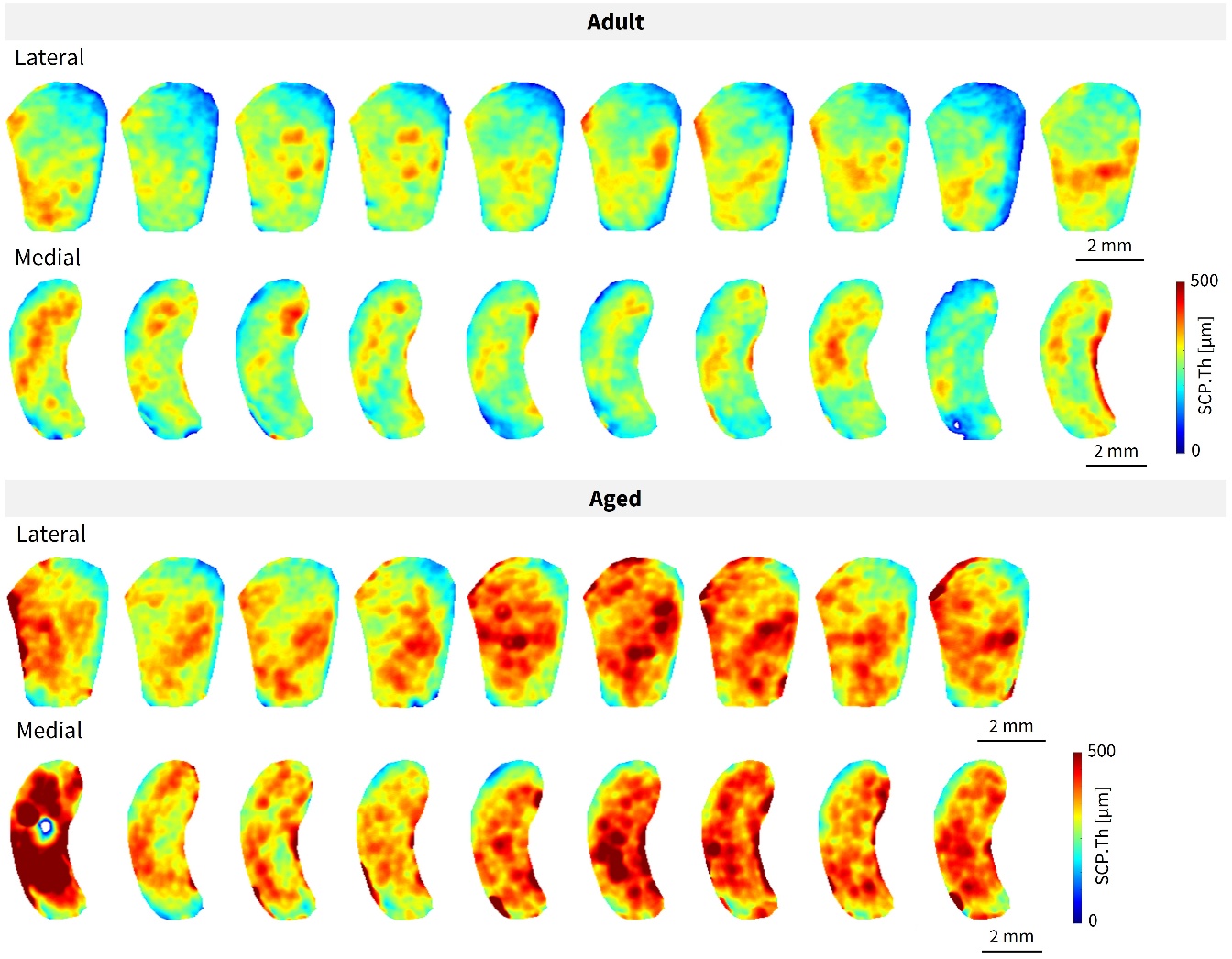

Figure S4: 2D projections of the subchondral plate thickness (SCP.Th) across all samples, revealing distinct thickness patterns in the medial and lateral plateaus. The lateral plateau exhibits a gradual thickness increase from the lateral (right) to the medial (left) sides of the plateau, with local thickening in the central-posterior region. The medial plateau presents a thick area forming a central band, as well as a small thick region at the very lateral (right) side.

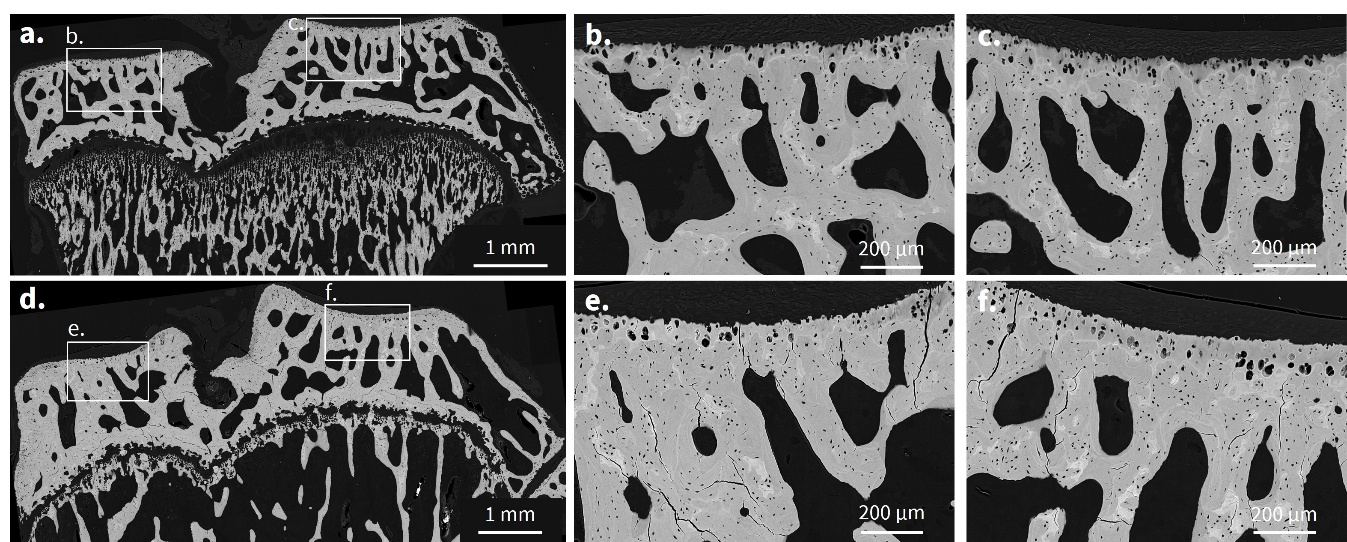

Figure S5: (a) qBEI images of a mid-frontal section from an adult sample, with (b, c) showing magnified views of the boxed regions. (d) qBEI image of a mid-frontal section from an aged sample, with (e, f) showing magnified views of the boxed regions, highlighting numerous cracks.

|  | **Tissue** | **Adult** | **Aged** | **Age** | **Location** | **Interaction** |
| --- | --- | --- | --- | --- | --- | --- |
| ${Ca}_{Mean}$ [wt %] | Bone | 25.12 ± 0.40 | 26.54 ± 0.39 | **<0.001** | **<0.001** | **0.0013** |
|  | Mineralized Cartilage | 22.71 ± 0.57 | 25.11 ± 0.54 |  |  |  |
| ${Ca}_{Width}$ [wt %] | Bone | 25.14 ± 0.43 | 26.60 ± 0.38 | 0.011 | **<0.001** | 0.570 |
|  | Mineralized Cartilage | 23.12 ± 0.33 | 25.23 ± 0.53 |  |  |  |
| ${Ca}_{Peak}$ [wt %] | Bone | 3.30 ± 0.23 | 3.05 ± 0.19 | **<0.001** | **<0.001** | **0.032** |
|  | Mineralized Cartilage | 5.50 ± 0.59 | 5.38 ± 0.59 |  |  |  |

Table S2: Mean ± SD values of parameters characterizing the mineralization frequency distribution in mineralized cartilage and subchondral bone for adult and aged animals. Results of two-way ANOVA are reported (p-values) considering age, location and their interaction.

| Parameter | Adult | Aged |
| --- | --- | --- |
| α [wt %] | 20.93 | 24.09 |
| β [µ$m^{-1}$] | 1.16 | 2.74 |
| dist_0_ [µm] | 1.68 | 1.47 |
| Ca_0_ [wt %] | 0.72 | 0.47 |
| TW [µm] | 9.13 | 3.86 |

Table S3: Parameters of the sigmoid function fitting the ummineralized to mineralized cartilage transition region: $\text{Ca }\left( \text{dist} \right)\text{ = }\text{Ca}_{\text{0 }}\text{+}\frac{\text{α}}{\text{1+}\text{e}^{\text{-β(dist-dist0)}}}$. The transition width (TW) is defined as: $\text{TW = }\text{dist}\left( \text{ Ca = 0.995*α+ }\text{Ca}_{\text{0}} \right) \text{-} \text{dist}\left( \text{ Ca = 0.005*α+ }\text{Ca}_{\text{0}} \right)$

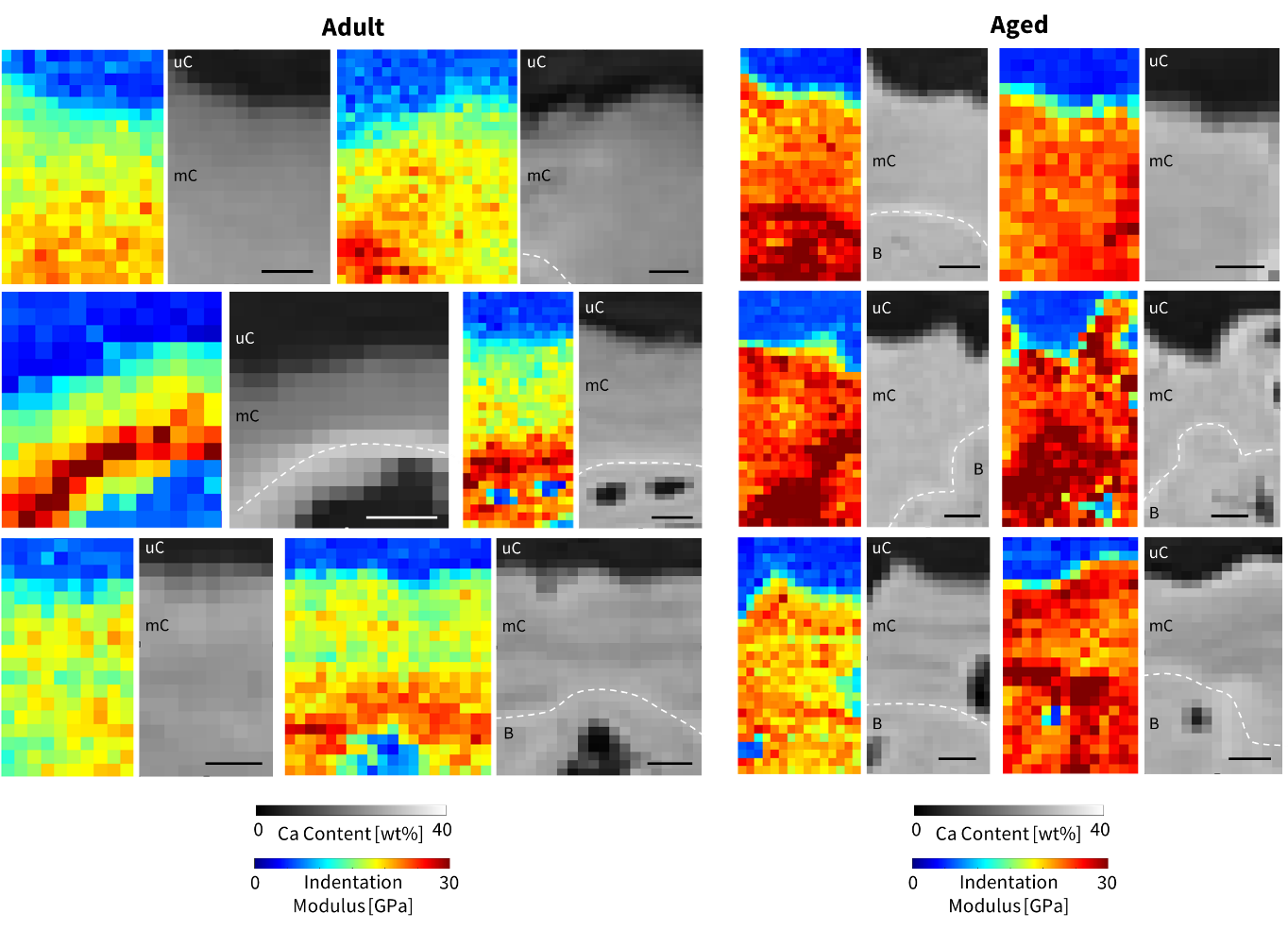

Figure S7: 2D maps of the indentation modulus on adult and aged samples and corresponding qBEI images. The osteochondral cement line is marked by white dotted lines. uC: unmineralized cartilage, mC: mineralized cartilage, SB: subchondral bone. Scale bar = 7µm.

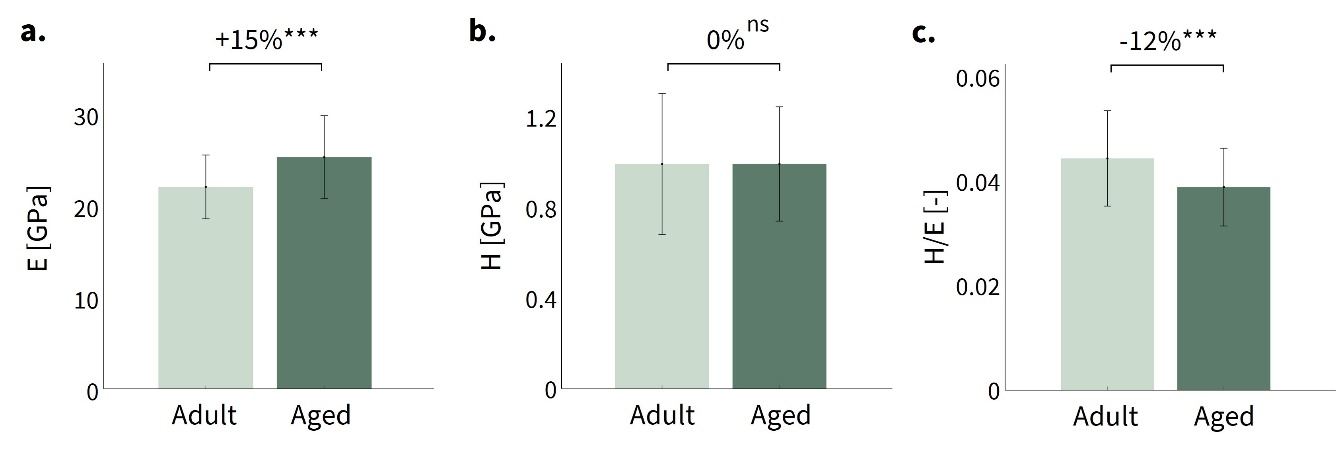

Figure S8: Quantitative comparison of (c) indentation modulus E, (d) hardness H, and (e) hardness/modulus ratio of subchondral bone for adult and aged rats. Data shown as mean (histogram height) with error bars representing one standard deviation. Statistical significance between data sets is indicated as follows: ***p < 0.001 and ns (not significant).

**References**

[1]	M. Doube, M.M. Kłosowski, I. Arganda-Carreras, F.P. Cordelières, R.P. Dougherty, J.S. Jackson, B. Schmid, J.R. Hutchinson, S.J. Shefelbine, BoneJ: Free and extensible bone image analysis in ImageJ, Bone 47 (2010) 1076–1079. https://doi.org/10.1016/j.bone.2010.08.023.
